# Predicting the Evolutionary and Functional Landscapes of Viruses with a Unified Nucleotide-Protein Language Model: LucaVirus

**DOI:** 10.1101/2025.06.14.659722

**Authors:** Yuan-Fei Pan, Yong He, Yu-Qi Liu, Yong-Tao Shan, Shu-Ning Liu, Jia-Hao Ma, Xue Liu, Xiaoyun Pan, Yinqi Bai, Zan Xu, Tingjun Hou, Zheng Wang, Jieping Ye, Jianguo He, Edward C. Holmes, Bo Li, Yao-Qing Chen, Zhao-Rong Li, Mang Shi

## Abstract

Predicting viral evolution and function remains a central challenge in biology, hindered by high sequence divergence and limited knowledge compared to cellular organisms. Here, we introduce LucaVirus, a multi-modal foundation model for viruses, trained on 25.4 billion nucleotide and amino acid tokens covering nearly all known viruses. LucaVirus learns biologically meaningful representations capturing relationships between sequences, protein/gene homology, and evolutionary divergence. Using these embeddings, we developed downstream models that address key virology tasks: identifying hidden viruses in genomic “dark matter”, annotating enzymatic activities of uncharacterized proteins, predicting viral evolvability, and identifying antibody candidates for emerging viruses. LucaVirus achieves state-of-the-art results in three tasks and matches leading models in the fourth with one-third the parameters. Together, these findings demonstrate the power of a unified foundation model to comprehensively decode the viral world and establish LucaVirus as an efficient and versatile platform for AI-driven virology, from virus discovery to functional and therapeutic predictions.

## Introduction

The global expansion of metagenomic sequencing has revealed an extraordinary and rapidly growing diversity of viruses across animals (*1*), plants (*2*) and a wide range of environments (*3, 4*). Despite their often small genomes, viruses encode nearly all biologically relevant information required for host interaction (*5*), immune evasion (*6*), and transmission (*7*)—making viral genomes and proteomes compact yet comprehensive blueprint of life. However, translating this genomic wealth into biological insight remains challenging (*8*). This bottleneck is mainly driven by the exceptionally high evolutionary rates of viruses (*9*), which leads to high levels of extreme sequence divergence (*10*), particularly among RNA viruses. Consequently, conventional bioinformatic methods based on sequence homology and conserved domain detection often lack the sensitivity and precision required to bridge the gap between viral discovery, functional characterization, and public health applications.

Recent advances in artificial intelligence, particularly large language models (LLMs), offer new opportunities to learn complex biological representations directly from sequence data. Models such as the ESM series (*11, 12*) and the Evo series (*13, 14*) have shown remarkable capabilities in protein function/structure prediction and *de novo* generation across cellular organisms. However, their application to virology has been constrained by a unique intersection of technical and biosafety challenges. First, viral sequences are substantially underrepresented in generalist foundation models trained predominantly on cellular organisms. Viral genomes exhibit distinct evolutionary constraints and composition biases that differ fundamentally from those in cellular life (*15*), including high coding density and widespread use of non-canonical mechanisms such as polyprotein processing and programmed ribosomal frameshifting (*16, 17*), features that are underrepresented in the data sets of existing models. Second, most existing models are unimodal, focusing exclusively on either protein or nucleotide sequences. Viral genomes, however, encode rich evolutionary information (*18, 19*)—such as codon usage biases indicative of host range—that is not explicitly captured in amino acid sequences. Conversely, proteins often preserve functional conservation better than rapidly evolving nucleotides, making protein-level representations indispensable for detecting remote homology. Finally, the rise of generative AI has introduced a biosecurity paradox: while models like Evo possess the architecture to learn genomic grammar, concerns regarding the potential synthesis of hazardous pathogens have led to the intentional exclusion or masking of viral sequences from their training corpora. This safety-driven blind spot leaves the virology community without high-capacity foundation models tailored to the full complexity of the virosphere.

To address these challenges, we present LucaVirus, a unified genome-protein language model specifically designed for viruses. Initialized from the general-purpose foundation model LucaOne (*20*), LucaVirus employs transfer learning followed by pre-training on one of the most comprehensive viral sequence collections assembled to date. By jointly modeling nucleotide and amino acid sequences, LucaVirus captures the unique grammatical and semantic dependencies underlying viral evolution, a design validated by rigorous ablation analyses. This unified framework supports diverse downstream applications, including discovery of highly divergent viruses, functional annotation, fitness landscape modeling, and antibody–antigen interaction prediction. Together, LucaVirus offers a generalizable and interpretable AI framework for advancing both basic virological research and pandemic preparedness.

## Results

### LucaVirus: a Unified Model for Viral Genomes and Proteins

LucaVirus is a unified genome-protein language model with one billion parameters, designed to interpret viral sequence at single-nucleotide and single-amino acid resolution (Fig. 1A). With the rapid expansion of viral diversity, there is an urgent need for a comprehensive and scalable model capable of capturing both genetic and functional characteristics across the entire virosphere. LucaVirus addresses this need by integrating nearly all known viral sequences into a single, cohesive framework.

**Figure 1.**
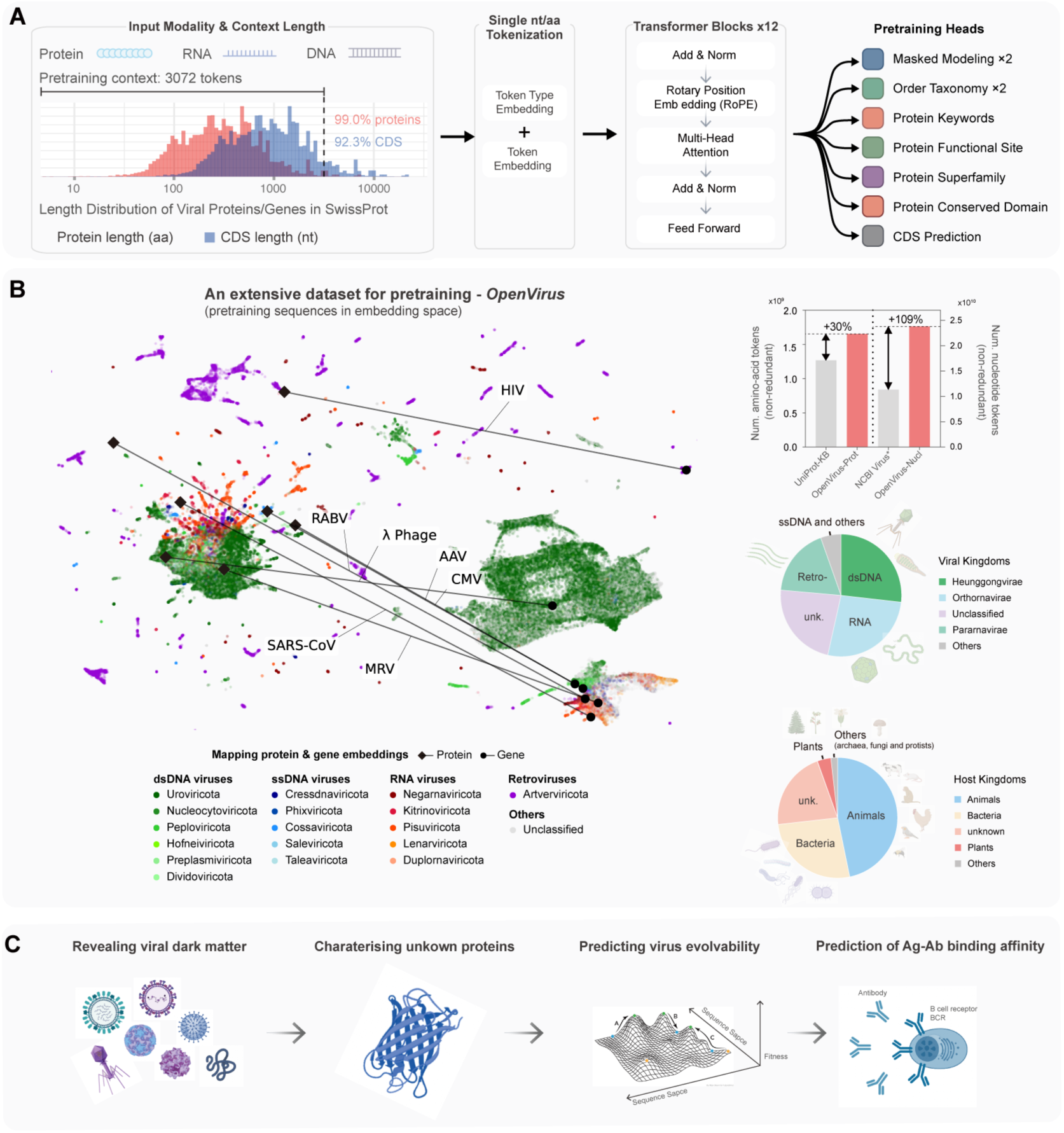
LucaVirus: A unified multi-modal foundation model for the global virosphere. **(A)** Architecture and pretraining tasks of LucaVirus. **(B)** Overview of the OpenVirus pretraining data set. Left: visualization of pretraining protein and nucleotide sequences in LucaVirus embedding space, with color indicating taxonomy and lines indicating the mapping between corresponding protein and nucleotide sequences. Right, bar plot: total tokens of non-redundant protein and nucleotide sequences in our pre-training data set compared with mainstream public databases – UniProtKB and NCBI Virus. Right, pie plot: taxonomic distribution and Host diversity within the training corpus. **(C)** Schematic overview of the four downstream applications enabled by LucaVirus.

To support such broad representation, we constructed OpenVirus, a large-scale pre-training corpus comprising 15.7 million viral sequences, including 10.4 million nucleotide and 5.2 million protein sequences, totaling 25.4 billion tokens (Fig. 1B). This data set substantially exceeds existing repositories, containing nearly twice the number of non-redundant tokens found in NCBI Virus and 30% more than UniProt (*21*). Importantly, OpenVirus incorporates recent large-scale viral discovery efforts (*10, 22–26*), contributing 1,663,163 previously uncatalogued “species-level” taxa (80% ANI) absent from public databases (Table S1). The data set spans all major viral realms—including double-stranded DNA viruses (*Heunggongvirae*, 27%), RNA viruses (*Orthornavirae*, 26%), and retroviruses (*Pararnavirae*, 20%), while covering viruses infecting hosts across all six cellular kingdoms (Fig. 1B).

To effectively learn from this diverse data and to prioritize sequence understanding over generation in light of biosecurity considerations, LucaVirus employs a customized encoder-only Transformer architecture with 1 billion parameters. The model was initialized with weights from LucaOne (*20*), a general-purpose biological foundation model. This transfer learning strategy allows LucaVirus to inherit fundamental biological sequence priors before extensive virus-specific training. To balance efficiency and expressiveness, we reduced the model to 12 transformer layers while expanding the context window to 3,072 tokens. This extended window is critical for viral genomics, allowing LucaVirus to capture long-range dependencies across 99% of viral proteins and 92% of coding sequences (CDS) in SwissProt (*21*) (Fig. 1A). Furthermore, LucaVirus was trained using a multi-task semi-supervised objective (*27, 28*) rather than standard masked language modeling, explicitly enforcing sequence-function associations and enhancing the model’s ability to decode viral functional semantics.

### Ablation Studies Demonstrate the Necessity and Robustness of a Unified Multi-Modal Design

To justify our architectural design, we benchmarked the standard LucaVirus against three variants: protein-only (LucaVirus-Prot), nucleotide-only (LucaVirus-Nucl), and a model trained exclusively on the masked language modeling (MLM) objective without biological supervision (LucaVirus-Mask). Across nine downstream tasks, the unified multi-modal LucaVirus outperformed all variants in seven, providing strong evidence that integrating nucleotide and protein representations is essential for a comprehensive understanding of viral biology (Fig. S2; Table S3). Although LucaVirus-Prot showed marginal advantages in capsid and enzyme prediction (by 0.02% and 0.22%, respectively), these gains were task-specific and did not generalize. And LucaVirus-Mask consistently underperformed across benchmarks, highlighting the limitations of purely syntactic language modeling and underscoring the importance of biologically informed training objectives for learning functional viral representations.

To further assess model stability and rule out “catastrophic forgetting” (*29*), we re-evaluated LucaVirus on ten general biological benchmarks originally developed for the LucaOne model(*20*). LucaVirus achieved comparable performance on 90% of these tasks (Fig. S3; Table S4), confirming that our transfer learning strategy effectively specializes the model for the virosphere while preserving fundamental biological syntax and generalization capacity.

### LucaVirus Recovers Fundamental Genomic Syntax and Non-canonical Translation Logic from Raw Sequence

LucaVirus maps viral sequences into a biologically structured, high-dimensional latent space without explicit supervision. Strikingly, nucleotide-level embeddings spontaneously recapitulate the fundamental rules of the central dogma, resolving multiple layers of genomic organization, including base identity, coding versus non-coding regions, codon phase structure, and synonymous codons (Fig. S4). This latent organization was quantitatively validated across all four features via permutational multivariate analysis of variant (PERMANOVA, 999 permutations, p < 0.05; Table S5), confirming that these biologically meaningful dimensions emerge intrinsically from model training.

To assess the linear separability of these representations, we conducted systematic linear probing benchmarks against state-of-the-art genomic language models (Fig. S5; Table S6) LucaVirus achieved 86%-100% accuracy across five core genomic discrimination tasks, including CDS identification, base identity, codon phase, amino acid identity, and codon identity. This level of performance was matched only by LucaOne (66%-100%). Notably, LucaVirus reached 93.8% accuracy in CDS identification, substantially outperforming state-of-the-art genome language models such as Evo-1-8k-base (62.9%) (*14*), Evo2-7B (74.3%) (*13*) and Nucleotide Transformer-v2-500M (74.7%) (*30*).

Beyond standard syntax, LucaVirus captures subtle, virus-enriched genomic features that are typically difficult to model. The embeddings precisely reconstruct relative nucleotide positions within codons (Fig. S6) and preserve phase consistency across programmed ribosomal frameshifting (PRF) boundaries. Quantitative evaluation using the FSDB database (*31*) (Fig. S7; Table S6) demonstrated that LucaVirus distinguishes codon phases with 98% precision, highlighting its exceptional sensitivity to non-canonical translational rules that are central to viral genome organization.

### Embedding Geometry Encodes Evolutionary Distance and Enables Embedding-Based Sequence Alignment

The geometric structure of the LucaVirus embedding space provides a high-fidelity representation of evolutionary divergence. Across 5,566 reference protein families and 938 “dark matter” families, we observed a strong and consistent correlation between embedding cosine similarity and genetic distance measured by p-distance (Fig. S8; Table S7). Importantly, this relationship generalizes well to sequences unseen during pre-training (Pearson’s r = 0.71; Fig. 2A) and extends robustly to nucleotide sequences (Table S8), demonstrating that LucaVirus embeddings capture fundamental evolutionary signals rather than data-set-specific artifacts.

**Figure 2.**
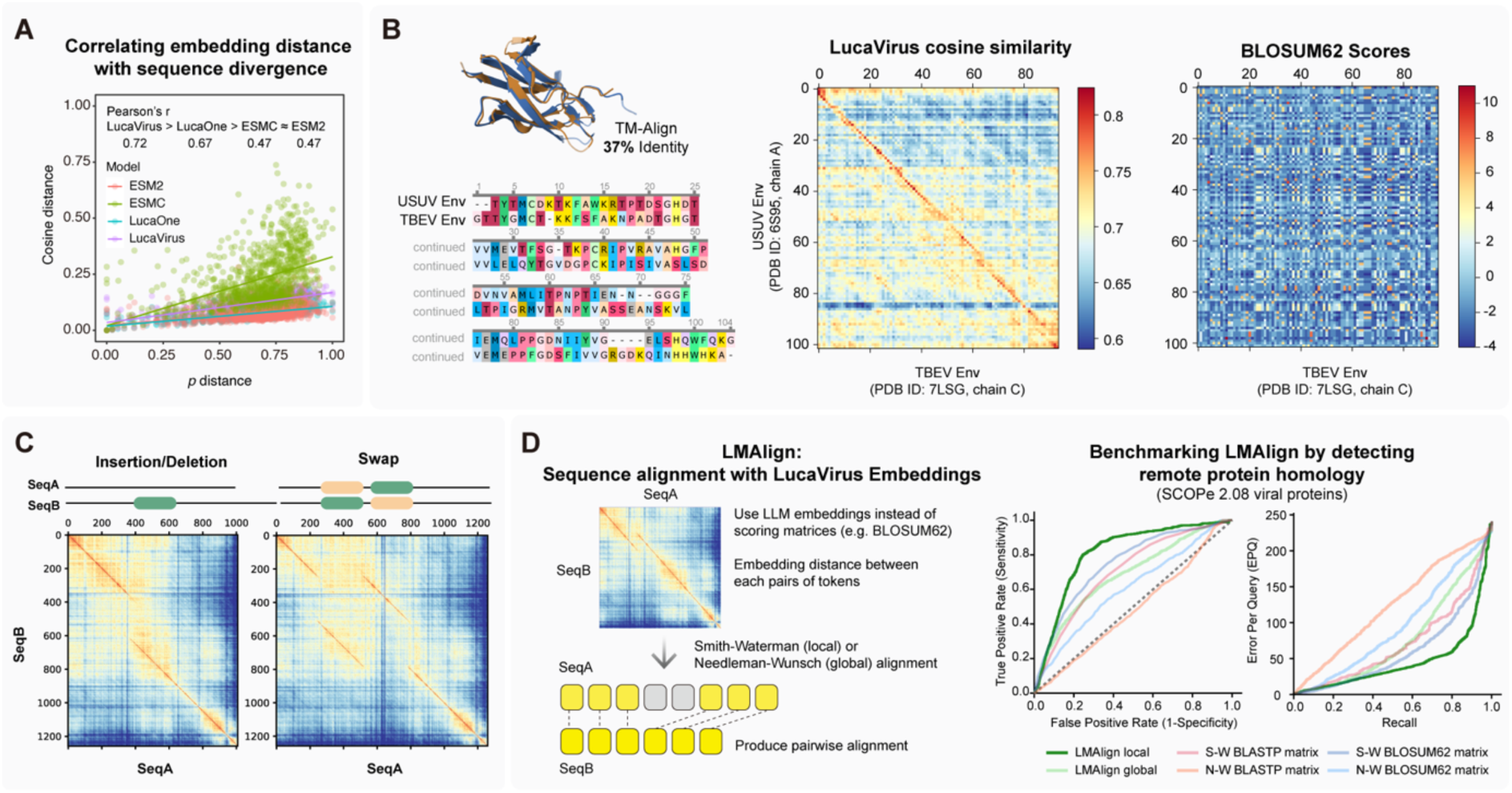
Learned representations by LucaVirus capture evolutionary divergence and enable embedding-based alignment. **(A)** Correlation between embedding space geometry and genetic distance. The scatter plot compares the Pearson’s *r* of cosine similarity against p-distance for LucaVirus versus other protein language models (LucaOne, ESMC, ESM2) on unseen viral protein families during LucaVirus pretraining. **(B)** Structural superposition (top) and sequence alignment (bottom) of two remote homologs and comparison of pairwise similarity matrices generated by LucaVirus embeddings versus the BLOSUM62 scoring matrix. **(C)** Visualization of structural variations in the embedding space. **(D)** Schematic of *LMAlign* alignment algorithm and benchmarks on remote viral protein homology detection. ROC curves and Error Per Query (EPQ) plots.

This representational capacity effectively captures remote homology. For example, in the envelope proteins of Usutu virus (USUV) and tick-borne encephalitis virus (TBEV), which share only 37% identity (Fig. 2B). Visualisation of pairwise residue scores demonstrate that LucaVirus embeddings distinctively highlight alignable regions, in sharp contrast to the noisy background produced by the standard BLOSUM62 scoring matrix (*32*)(Fig. 2B). The embeddings further exhibit sensitivity to structural variations such as indels and domain rearrangements (Fig. 2C), indicating that contextual and structural constraints are implicitly encoded in the latent space. Building on these insights, we developed LMAlign, a novel sequence embedding-based sequence alignment algorithm that leverages LucaVirus representations to guide alignment (Fig. 2D; Fig. S9). Benchmarking on the SCOPe 2.08 data set (*33*) confirmed that LMAlign significantly outperforms classic Smith-Waterman and Needleman-Wunsch algorithms in detecting remote structural homology (Fig. 2D).

### LucaVirus Helps Resolves Key Challenges in Virology

Decoding the global virosphere requires overcoming a series of interconnected barriers that span from virus discovery to pandemic preparedness. Beyond the fundamental challenge of identifying highly divergent viruses hidden within genomic dark matter, researchers face the critical bottleneck of assigning biological functions to the vast majority of viral proteins that lack cellular homologs. Furthermore, effective viral surveillance demands not only static characterization but also the dynamic forecasting of viral fitness landscapes and the rapid identification of neutralizing antibodies. To demonstrate the versatility of LucaVirus, we systematically assessed its performance across these four pivotal domains, utilizing lightweight downstream models to interpret the rich biological semantics encoded in its pre-trained embeddings.

For each task, we employed lightweight, task-specific downstream models that operate directly on the embeddings generated by pre-trained language models (see Methods). These downstream models share a unified, minimal architecture comprising a single attention-based pooling layer, a fully connected layer, and a task-specific prediction head (Fig. S10), while keeping the upstream language model frozen. This design minimizes trainable parameters, improves computational efficiency, and reduces dependence on large-scale, labeled data sets—a major advantage in virology, where experimentally validated data are scarce.

### Challenge 1: High-sensitivity Discovery of Viral Hallmark Proteins from Genomic Dark Matter

Building on our previous work (*10*), we targeted two hallmark viral proteins—the RNA-dependent RNA polymerase (RdRP) and viral capsid proteins—to uncover unclassified RNA viruses present within genomic dark matter (Fig. 3A). RdRP is a highly conserved enzyme critical for RNA virus replication and rarely found in cellular organisms (*34*), making it a robust marker for virus discovery (*35*). In contrast, viral capsid proteins are structurally essential but highly divergent, posing a major challenge for conventional homology-based detection (*36*).

**Figure 3.**
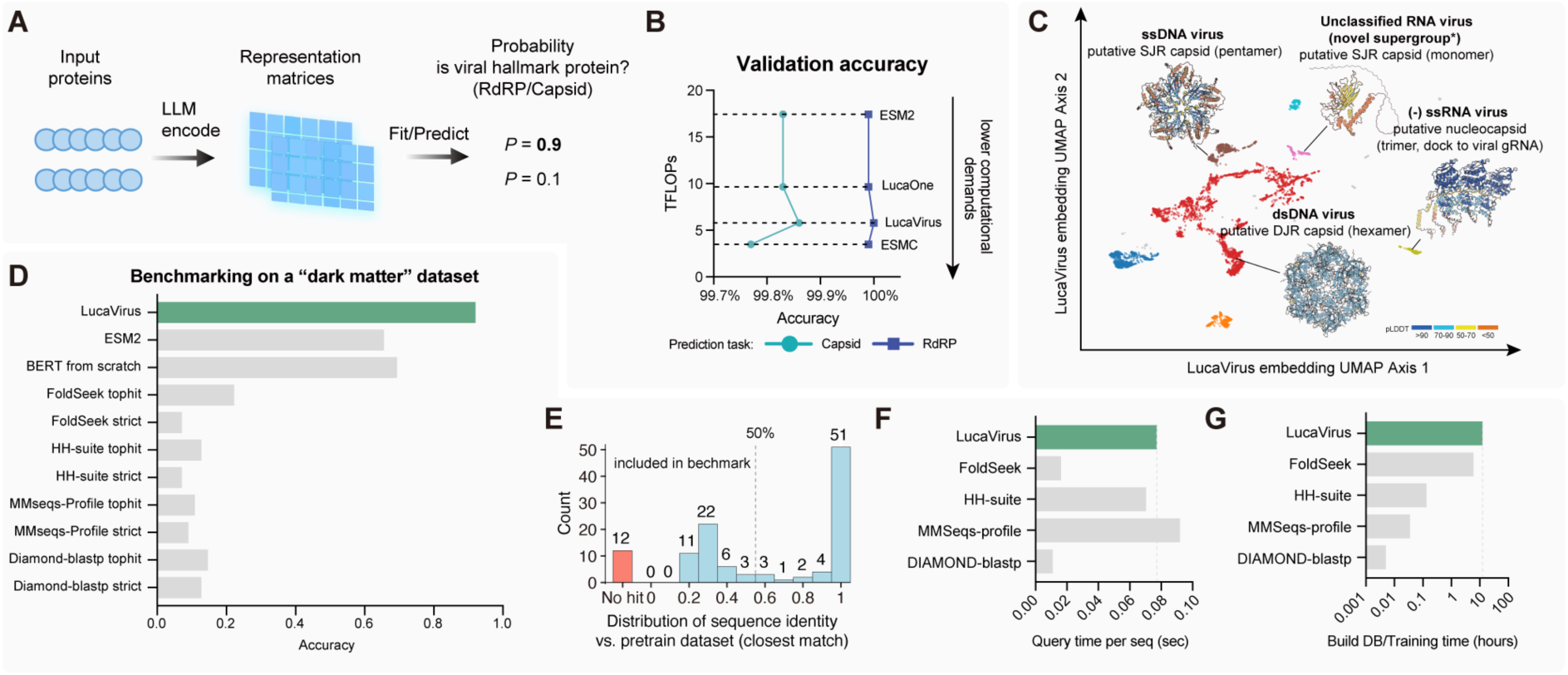
Uncovering viral dark matter through efficient hallmark protein identification. **(A)** Workflow for the binary classification of viral hallmark proteins (RdRP and Capsid) using LucaVirus embeddings. **(B)** Comparison of the accuracy and computational complexity (floating point operations per inference, TFLOPs) among three large language models in RNA-dependent RNA polymerase (RdRp) and capsid protein prediction. **(C)** UMAP visualization of the LucaVirus embedding space for viral capsid proteins. **(D)** Performance on a “dark matter” test set of divergent viral proteins. **(E)** Distribution of sequence identity between the “dark matter” test set and the nearest matches in the pre-training data set. **(F–G)** Efficiency comparison for query time per sequence **(F)** and database construction/training time **(G)**.

On a validation set comprising 23,000 RdRPs and 48,000 capsid proteins, the LucaVirus hallmark protein identification model achieved accuracies of 100% for RdRP and 99.86% for capsids (Fig. 3B; Table S6-7). Notably, LucaVirus matched the performance of the substantially larger ESM2-3B model while reducing reference cost from 17 TFLOPs to 5 TFLOPs per inference, demonstrating markedly improved efficiency. Through visualization, we showed that LucaVirus robustly generalizes across diverse capsid architectures, including nucleocapsids, single jelly roll (SJR), and double jelly roll (DJR) folds, despite substantial sequence and structural heterogeneity.

To rigorously evaluate real-world generalization, we benchmarked LucaVirus on a curated set of 115 structurally validated proteins from recently discovered RNA viruses (68 capsids and 47 non-capsids; Extended Data Set 1) (*10*), after excluding any sequences sharing >50% identity with the pre-training data. Under these stringent conditions, LucaVirus achieved 100% recall and accuracy (Fig. 3D; Table S9). In contrast, ESM2-3B reached only 66% accuracy (65.3% recall), while structure-based (FoldSeek) (*37*) and sequence- or profile-based methods (DIAMOND blastp (*38*), HHblits (*39*) and MMseqs2 (*40*)) showed accuracies below 22.6% and recalls below 10% (Fig. 3D).

Finally, we assessed scalability on large-scale data sets using identical hardware (128-core, 256-threads CPU, single NVIDIA A100 GPU). LucaVirus achieved a per-sequence inference time of 0.077 seconds, positioning it between HHblits and MMseqs2 in speed (Fig. 3F). This inference speed makes LucaVirus the only method that combines high sensitivity with the throughput required for large-scale metagenomic mining. Although model training requires a one-time computational cost comparable to database indexing for FoldSeek (Fig. 3G), LucaVirus thereafter provides an efficient and scalable solution for systematic exploration of viral genomic dark matter.

### Challenge 2: Breaking the Functional Annotation Bottleneck of the Viral Proteome

Although metagenomics has rapidly expanded the known virosphere, functional annotation of viral proteins remains a critical bottleneck due to extreme sequence divergence, which severely limits conventional homology-based methods. We therefore evaluated LucaVirus for viral protein function prediction across increasingly challenging benchmarks.

We first evaluated LucaVirus on a standard viral Enzyme Commission (EC) number prediction task using UniProtKB-derived data sets (Fig. 4A). LucaVirus achieved 99.97% accuracy with an F1 score of 0.9889 (Fig. 4B; Table S10), matching the performance of the much larger ESM2-3B model while operating with substantially fewer parameters, demonstrating its efficiency.

**Figure 4.**
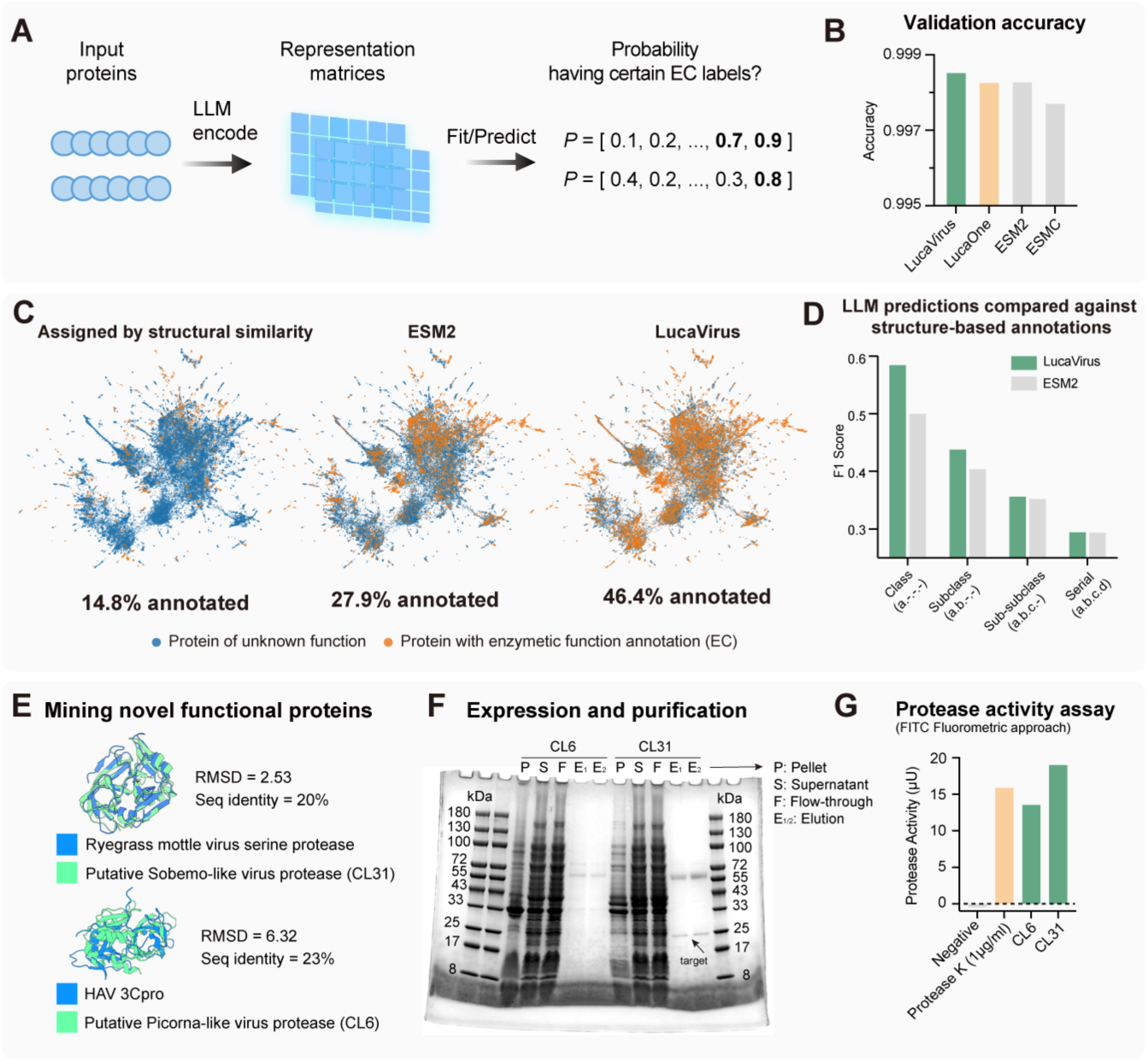
Functional annotation and experimental validation of novel viral enzymes. **(A)** Schematic of the enzyme function prediction task, mapping input sequences to Enzyme Commission (EC) probabilities. **(B)** Validation accuracy comparison across models. LucaVirus achieves superior accuracy (>0.998) in assigning EC labels to viral proteins. **(C)** UMAP visualization of the protein embedding space from the Nomburg et al. (2024) data set (*8*). Points are colored by predicted (orange) or unknown (blue) enzymatic function. **(D)** Performance evaluation against structural ground truth on the Nomburg et al. (2024) data set (*8*). **(E)** Discovery of two novel viral protease and structure alignment against known protease. **(F)** SDS-PAGE analysis of the expression and purification of two putative viral proteases (CL6 and CL31) predicted by the model. **(G)** *In vitro* proteolytic activity assay using a fluorescence-based method.

Because standard UniProt benchmarks may be saturated—making it difficult to differentiate model performance—we next evaluated LucaVirus on a rigorously curated, structure-based data set (*8*) in which viral proteins were annotated through structural alignment. While the original structural approach annotated only 14.8% of the proteins and the ESM2-based model reached 27 (*20*).9%, LucaVirus expanded annotation coverage to 46.4% (Fig. 4C). When benchmarked against structural ground truth—excluding sequences with >50% identity to pretraining data—LucaVirus achieved a recall of 44.5% and an F1 score of 0.587 at the enzyme class level, outperforming ESM2-3B (recall: 35.1%, F1: 0.5; Table S10).

To test real-world discovery potential, we applied LucaVirus to identify putative novel proteases (EC 3.4.-.-) from the genomic “dark matter” of recently discovered RNA viruses (*10*). LucaVirus identified 8,074 candidate proteins (Extended Data Set 2), from which 46 clusters were selected for structural validation using ColabFold (*41*) (Extended Data Set 3). Eight clusters displayed clear structural similarity to known proteases (Foldseek E-value < 10^-5^), despite sharing only ∼12% sequence identity, well within the homology “twilight zone”. Two candidates (CL31 and CL6) were experimentally validated: both were successfully expressed using an *E. coli* system, showed oligomerization consistent with viral proteases, and exhibited protease activity *in vitro* (Fig. 4G). Remarkably, these discoveries were achieved using sequence data alone without structural inputs, which underscores LucaVirus’s ability to infer deep sequence-to-function relationships beyond the reach of conventional alignment- and structure-based methods.

### Challenge 3: Predicting the Viral Fitness Landscapes and Evolutionary Potential

Accurately modeling viral fitness landscape is essential for predicting viral evolvability. Unlike cross-species functional annotation, this task requires resolving subtle fitness effects of intra-species variation. We evaluated LucaVirus using deep mutational scanning (DMS) data, which systematically quantify the fitness impact of single mutations (*42*). Focusing first on receptor-binding affinity of SARS-CoV-2 Spike RBD variants (Fig. 5A-B; Table S11), LucaVirus achieved state-of-the-art performance with a Spearman correlation of 0.93, outperforming generalist models including LucaOne (0.90), ESM2-3B (0.84), and ESMC-600M (0.71) (Fig. 5C). Notably, LucaVirus retained strong performance using nucleotide sequences as input (Spearman = 0.87), surpassing genome language models like Evo2-7B (*13*), DNABert2 (*43*) and Nucleotide Transformer-2.5B-multi-species (*30*) (Fig. 5C; Table S11), and demonstrating robust cross-modal consistency. These models were trained on early SARS-CoV-2 variants and tested on later, unseen variants, highlighting LucaVirus’s ability to extrapolate from limited experimental data to emerging strains.

**Figure 5.**
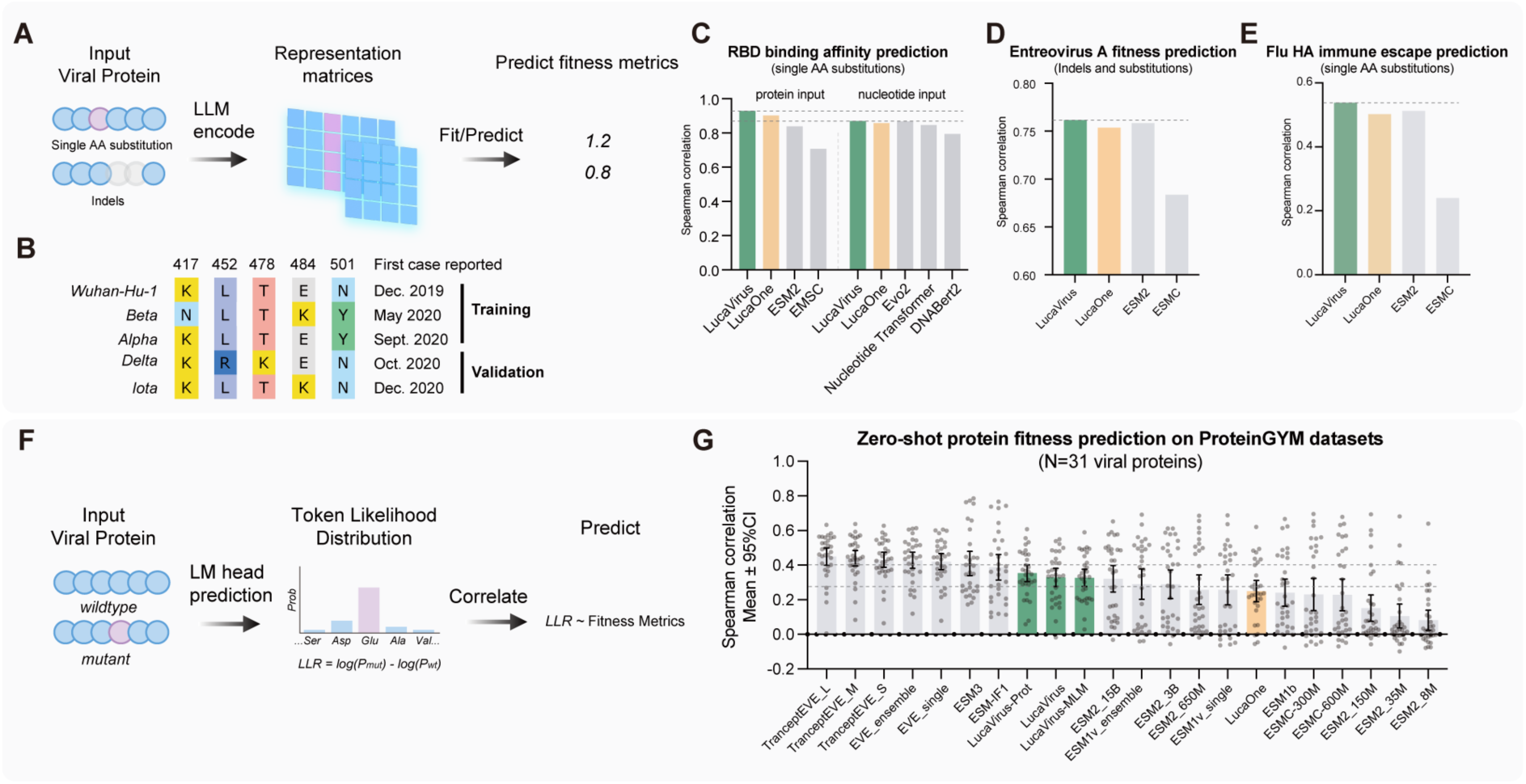
Forecasting viral evolvability and fitness landscapes via zero-shot inference. **(A)** Workflow for predicting fitness effects of single amino acid substitutions and indels in using a supervised approach. **(B)** The temporal split of SARS-CoV-2 variants used for training and validation. **(C–E)** Spearman correlation coefficients between model predictions and experimental Deep Mutational Scanning (DMS) data for SARS-CoV-2 RBD binding affinity **(C)**, Enterovirus A fitness **(D)**, and Influenza Hemagglutinin (HA) immune escape **(E)**. **(F)** Schematic of the zero-shot prediction strategy using the Log-Likelihood Ratio (LLR) of mutant versus wildtype tokens. **(G)** Zero-shot performance benchmark on the ProteinGym data set. The scatter plot shows the mean Spearman correlation (with 95% CI) for LucaVirus compared to state-of-the-art models.

To assess generality beyond coronaviruses, we extended the evaluation to additional viral families, including Enterovirus polyprotein fitness (Fig. 5D) and the Influenza A Hemagglutinin (HA) immune escape landscapes (Fig. 5E). LucaVirus consistently achieved superior performance across both tasks, confirming its broad applicability across viral families.

Finally, we evaluated LucaVirus’s intrinsic fitness awareness using the zero-shot ProteinGYM benchmark (*44*). LucaVirus demonstrated competitive performance with state-of-the-art models—including the MSA-dependent EVE (*45*) and Tranception series models (*46*), and the 98B-parameter ESM3 (*12*)—despite operating without alignment constraints and with significantly fewer parameters. This result reflects expected architectural trade-offs: EVE explicitly models evolutionary history via multiple sequence alignments (MSAs), while ESM3 leverages a massive parameter space (98B vs. 1B for LucaVirus); however, these computationally intensive strategies still conferred no statistically significant advantage over LucaVirus (*t*-test, p>0.05).

### Challenge 4: Accurate Antibody-Antigen Interactions Modeling for Pandemic Readiness

LucaVirus also demonstrated strong capability in identifying antibodies from repertoires targeting emerging viruses (Fig. 6A). Evaluated on the CoV-AbDab coronavirus antibody database (*47*), LucaVirus achieved 93% accuracy in predicting binding between antibodies and the SARS-CoV-2 spike protein (Fig. 6B; Table S9), indicating robust baseline performance.

**Figure 6.**
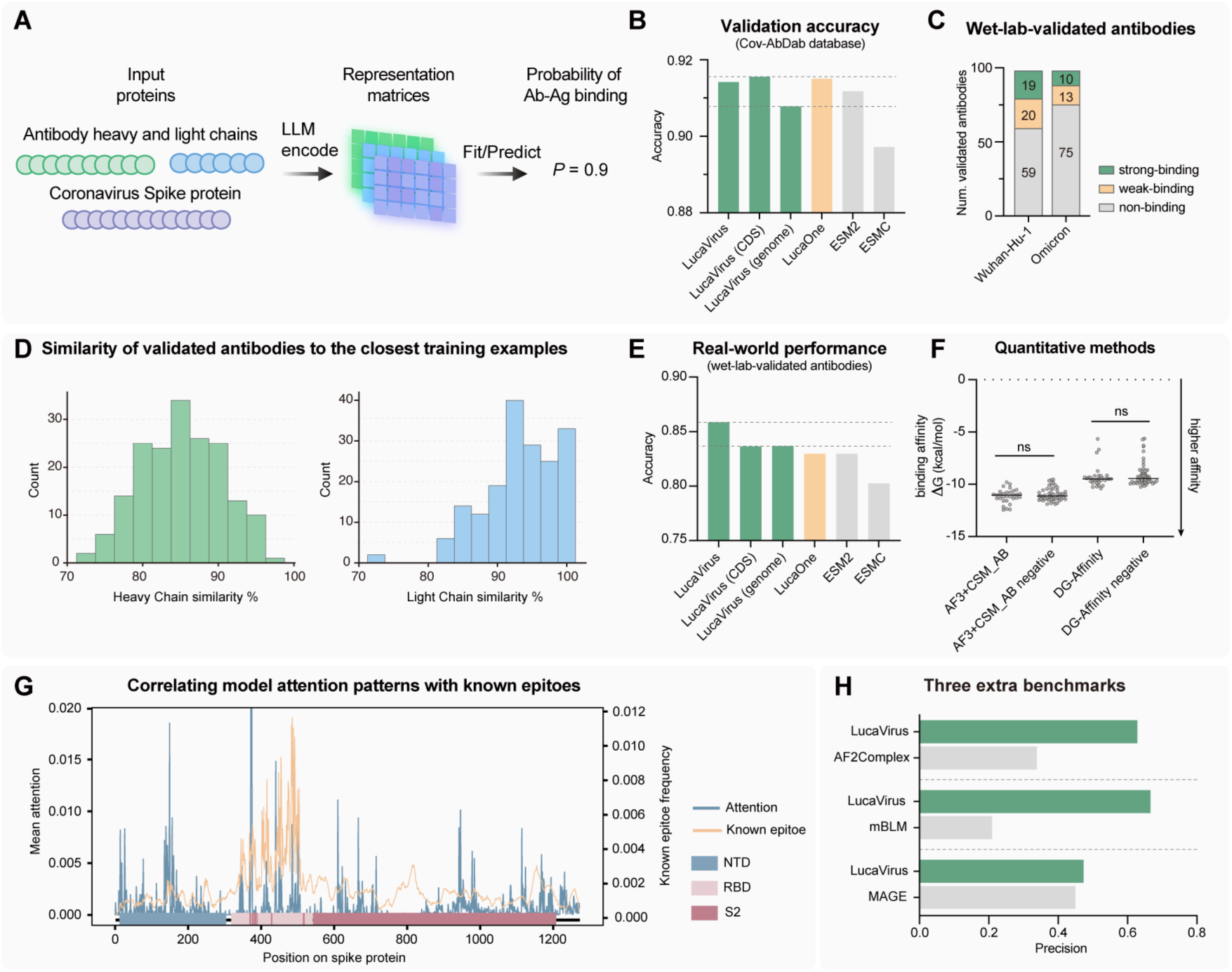
Accurate prediction of virus-specific antibody-antigen interactions. **(A)** Schematic diagram of the antibody-antigen binding prediction task. **(B)** Validation accuracy on the CoV-AbDab database. **(C)** Experimental validation using real-world antibody test data, illustrating the distribution of antibodies binding to the S protein of Wuhan-Hu-1 and Omicron strains in this data set. **(D)** Histograms showing the sequence similarity of validated antibodies to the closest examples in the training set. **(E)** Real-world performance accuracy on the wet-lab validated antibody set. **(F)** Predictions of two qualitative approaches on real-world data sets. **(G)** Attention analysis of the model. The line plot overlays the mean attention weights of the model onto the SARS-CoV-2 spike protein sequence. High attention regions (blue) correlate strongly with known epitopes (orange) in the N-terminal domain (NTD) and Receptor Binding Domain (RBD). **(H)** Benchmarking against three independent, state-of-the-art models. Precision (True Positive / All Predicted Positive) is used to indicate the model’s efficacy in identifying novel, true-positive binders.

To assess real-world utility, we simulated an antibody discovery pipeline using human B-cell receptor (BCR) repertoires. We randomly selected 98 antibodies from single B-cell sequencing of convalescent COVID-19 patients and experimentally measured their binding to both Wuhan-Hu-1 and Omicron spike proteins (Fig. 6C; Fig. S11, Table S12-13; Extended Data Set 4). After stringent filtering to exclude antibody with fewer than 10 amino acid mismatches from training data, 80 validated antibodies were retained for evaluation (Fig. 6D). On this data set, LucaVirus achieved accuracies of 84.6% (Wuhan-Hu-1) and 89.8% (Omicron), outperforming the protein language models tested (Fig. 6E) and structure-based methods—including AlphaFold 3 (*48*) combined with CSM-Ab (*49*) and DG-affinity (*50*) (Fig. 6F). Attention-weight visualization further revealed strong concordance between high-attention regions and known spike epitopes, suggesting that LucaVirus explicitly captures biologically relevant interaction sites (Fig. 6G).

Notably, LucaVirus exhibits unique cross-modal flexibility. When retrained to predict binding affinity using spike-coding CDS sequences or entire viral genomes instead of protein sequences, the model maintained high accuracy (both 84.3%; Fig. 6D; Table S9), demonstrating its ability to extract antigenic signals directly from raw genomic data and streamline discovery pipelines where protein annotation may be missing.

Finally, we benchmarked LucaVirus against three state-of-the-art models—mBLM (*51*), AF2Complex (*52*), and MAGE (*53*)—across three independent data sets (Fig. 6H). LucaVirus consistently outperformed these competitors, achieving precisions ranging from 47% to 66%. In practical terms, this translates to an experimental hit rate of approximately one in every two candidates, offering a highly efficient pipeline for therapeutic antibody screening and discovery.

Crucially, all benchmark sequences were stringently filtered to ensure at least 10 mismatches from the training set, underscoring the robustness and generalization capability of LucaVirus in antibody-antigen interaction prediction.

## Discussion

Modeling the evolutionary and functional landscape of viruses remains a fundamental challenge, largely due to their extensive sequence diversity and the limited biological understanding compared to cellular organisms. To address this gap, we present LucaVirus—the first unified, multi-modal foundation model purpose-built for virology. By integrating nearly all available viral sequence data across both nucleotide and protein modalities, LucaVirus establishes a comprehensive framework for interpreting viral biology at multiple levels.

Our results highlight the advantages of domain-specific foundation models as a powerful complement to general-purpose large language models. While generalist models offer broad biological coverage, their substantial computational costs and lack of domain granularity often constrain their practical deployment in specialized fields (*27, 54*). In our comprehensive benchmarking, the 1B-parameter LucaVirus consistently matched or outperformed a range of leading generalist models—including both protein-centric (e.g., the ESM series) and genome-centric (e.g., Evo, Nucleotide Transformer) architectures—across diverse virological tasks. This underscores that domain specialization can deliver substantial gains in both performance and parameter efficiency.

We further validate the effectiveness of transfer learning in biological sequence modeling. A central concern in fine-tuning is “catastrophic forgetting” (*29*), whereby specialization erodes general knowledge. By benchmarking LucaVirus on the original LucaOne’s ten general biological tasks (*20*), we show LucaVirus retains core biological syntax while gaining virus-specific expertise. This establishes a robust and generalizable paradigm: initializing from a broad biological foundation and subsequently refining with domain-specific pre-training offers an effective strategy for building specialized biological AI models.

Our ablation analyses provide insight into the architectural requirements for viral modeling. Both the unified nucleotide-protein representation and the semi-supervised, biologically informed pre-training objective proved essential for strong performance in supervised downstream tasks, consistent with prior studies (*20, 27, 28*). Interestingly, in zero-shot fitness prediction, the pure mask-prediction variant slightly outperformed the standard LucaVirus, reflecting fundamental trade-off. While mask-prediction objective optimizes perplexity and benefits zero-shot probability estimation, the semi-supervised strategy explicitly forces the model to learn sequence-function associations, yielding superior performance in biologically relevant supervised tasks.

This distinction underscores a key difference between biological sequences and natural language. Unlike natural language, which is largely self-descriptive (*55*), biological sequences encode function implicitly and context-dependently. Pure autoregression or masking may capture statistical patterns but fail to recover functional semantics. By incorporating explicit functional supervision, LucaVirus extracts biologically meaningful representations—capturing codon structure, synonymous mutations, and phylogenetic relationships—that form a stronger foundation for addressing complex, label-dependent virological challenges.

Furthermore, the architectural decision to employ a bidirectional encoder (BERT-style) rather than a unidirectional generative decoder (GPT-style) was strategic. While generative architectures (GPT-like) have demonstrated utility in de novo sequence design by optimizing likelihood estimation (*46, 56*), they rely on unidirectional causal masking which limits access to global contextual information. In contrast, our encoder-only architecture leverages bidirectional attention to capture the epistatic interactions and long-range dependencies essential for functional characterization (*11*). Crucially, this architecture serves a “safety-by-design” principle. By restricting the model to representation learning and analytical inference, we deliberately exclude the generative capabilities required for *de novo* pathogen design (*57*).

Beyond predictive accuracy, LucaVirus enables a paradigm shift in computational virology. Rather than serving solely as a predictive tool, it functions as a hypothesis-generating platform—an “in silico virology lab”. As demonstrated in evolvability and antigen-antibody interaction tasks, LucaVirus enables rapid, large-scale *in silico* exploration of mutational effects and candidate antigen-antibody pairs, allowing researchers to prioritize hypotheses before undertaking resource-intensive experimental assays. Similar frameworks have transformed fields such as neuroscience (*58, 59*), and LucaVirus now offers comparable potential for accelerating discoveries in virology.

Looking forward, LucaVirus points toward a new paradigm in virology, in which domain-specific foundation models become core infrastructure for understanding, monitoring, and responding to viral threats. By transforming fragmented viral sequence data into unified, interpretable representations, LucaVirus enables a shift from reactive analysis to proactive exploration of viral diversity, function, and evolution. As viral sequencing continues to accelerate, such models will be essential for translating raw data into actionable biological insight at scale. More broadly, LucaVirus demonstrates how biologically grounded, safety-aware AI systems can function as durable discovery platforms rather than task-specific tools. Future integration of structural, host, and ecological information may further enhance its ability to anticipate viral emergence and inform public health preparedness, positioning LucaVirus as a foundational framework for responsible innovation in virology.

## Materials and Methods

### Data Collection and Pre-processing

**The OpenVirus Corpus.** We curated OpenVirus, a comprehensive data set of viral sequences used to train the LucaVirus model. This data set comprises 15.7 million viral sequences, totaling 25.4 billion tokens—including 23.7 billion nucleotide tokens from 10.4 million sequences and 1.6 billion amino acid tokens from 5.2 million protein sequences. Nucleotide sequences were collected from NCBI Virus and seven large-scale virus discovery studies (*10, 22–26*) to ensure coverage of uncatalogued viral diversity. Protein sequences were obtained from UniProtKB (*60*) and the ColabFold environmental database (envdb) (*41*), which integrate and de-redundifies large environmental databases such as BFD and MGnify (*61*). To mitigate sampling bias and capture intra-species variation, nucleotide sequences were clustered using MMseqs2 (*40*) easy-cluster at 80% coverage and 80% ANI. Up to 10 non-redundant sequences were sampled per cluster, ensuring balanced representation across species-level taxa defined by 80% ANI. The final data set spans all known viral realms (including *Heunggongvirae*, *Orthornavirae*, and *Pararnavirae*) and encompasses virus-infecting hosts across all cellular domains.

**Biological Annotations.** To support biologically informed pre-training, we enriched the sequence data with functional and taxonomic annotations. For nucleotide sequences, annotations were primarily derived from GenBank GBFF files, from which coding region coordinates and order-level taxonomic labels were extracted. Sequences lacking annotations were retained with missing fields left blank. Protein annotations were obtained from UniProtKB (*60*) and InterPro (*62*), with InterPro providing details on functional sites, conserved domains, and homologous superfamilies. Taxonomic labels were defined at the order level based on 63 categories, with missing entries assigned a default label of −100. Functional annotations included: (i) 946 functional site categories (e.g. active sites, binding sites, conserved sites, and post-translational modifications), (ii) 3,460 homologous superfamilies, (iii) 1,659 conserved domains, and (iv) 603 functional keyword groups.

### Model Architecture and Pre-training

**Architecture.** LucaVirus is a 1-billion-parameter, encoder-only Transformer model purpose-built for viral sequence analysis. It features 12 transformer layers, a hidden dimension of 2,560, and an expanded context window of 3,072 tokens. We strategically adopted an encoder-only architecture rather than a generative framework for two reasons: (i) Interpretability: encoder models extract robust feature matrices rather than generating text, enabling direct interpretation and analysis of learned biological features; and (ii) Biosecurity: by design, encoder-only models lack generative capacity, thereby mitigating dual-use risks associated with *de novo* pathogen sequence generation.

**Model Scaling and Structural Adaptations.** We determined the model’s parameter size by adopting a parameter-to-token ratio of approximately 1:25 relative to our available training data. This yielded a 1B-parameter architecture well supported by the OpenVirus corpus, which contains twice the non-redundant tokens of NCBI Virus and 30% more than UniProtKB. To reduce parameter relative to the 1.8B LucaOne parent model (*20*) while preserving representational capacity, we retained the high-dimensional embedding size (2,560) but reduced the network depth (from 20 to 12 layers) and the prediction head size (from 40 to 20). To better capture long-range dependencies characteristics of viral genomes, we extended the maximum context length from 1,280 to 3,072 tokens. This expansion, optimized for NVIDIA A100 (80GB) hardware, accommodates the full-length sequences of 99.1% of full-length viral proteins in SwissProt (*60*).

**Transfer Learning and Pre-training Protocol.** LucaVirus was initialized from LucaOne checkpoint at step 1,760,000 (*20*) and subsequently refined using a semi-supervised pre-training strategy. This approach enables inheritance of broad biological sequence priors followed by virus-specific specialization. The pre-training process integrated self-supervised masked language modeling (MLM) with seven biologically relevant supervised tasks, including sequence-level classification (order-level taxonomy for nucleotides/protein sequences and functional keyword prediction) and token-level annotation (gene boundaries, homologous superfamilies, conserved domains, and functional sites). To balance these objectives, we assigned a loss weight of 1.0 for MLM and keyword prediction tasks, and 0.2 for the remaining auxiliary tasks.

Training was conducted on eight NVIDIA A100 GPUs (Alibaba Cloud) for 70 days, using a learning rate of 2×10^-4^ with warm-up phase, a batch size of 8, and gradient accumulation 32 steps. The model was trained for 3.8 million steps (approximately 1.9 epochs), processing over 49 billion tokens. Convergence was monitored by tracking average loss every 4,000 steps.

Benchmarking across all downstream tasks confirmed that both the unified nucleotide–protein representation and the semi-supervised training objectives are essential for optimal performance. In addition, evaluation on general biological benchmarks from the original LucaOne study verified that the transfer learning strategy effectively avoids catastrophic forgetting.

**Ablation Studies.** To validate our architectural and training strategies, we developed three ablation variants: LucaVirus-Prot (protein-only), LucaVirus-Nucl (nucleotide-only), and LucaVirus-Mask (masked language modeling only, without additional biological supervision). We benchmarked these variants against the standard LucaVirus model across all downstream tasks (detailed below).

To assess potential “catastrophic forgetting”—the erosion of fundamental biological capabilities during domain-specific pre-training—we compared LucaVirus against the original LucaOne foundation model (*20*). We employed the standardized evaluation suite established in the LucaOne study, which encompasses ten general biological tasks across three domains: (1) Fundamental Sequence-Function Mapping: We utilized the Central Dogma task to assess the model’s intrinsic ability to map DNA sequences to protein products without explicit alignment supervision. Additionally, we evaluated Gene Taxonomy Prediction (at Superkingdom, Species, and Genus levels) to verify the retention of phylogenetic signals. (2) Single-Molecule Property Prediction: To confirm the preservation of molecular feature recognition, we included Non-coding RNA Family classification (ncRNAFam) for RNA structural semantics, Prokaryotic Protein Subcellular Localization (ProtLoc) for sorting signal identification, and Protein Stability (ProtStab) regression for biophysical property estimation. (3) Molecular Interaction Prediction: To evaluate cross-modal and pairwise interaction capabilities, we tested Protein-Protein Interaction (PPI), ncRNA-Protein Interaction (ncRPI), and Influenza A Antigenic Relationship (InfA) prediction.

### Embedding Analyses

**Nucleotide Representation and Biological Syntax.** To visualize the nucleotide-level representations learned by LucaVirus, we extracted embedding matrices from 30 annotated Enterovirus reference genomes obtained from NCBI/GenBank. Dimensionality reduction was performed using Principal Component Analysis (PCA). To assess biological interpretability, embeddings were colored by four attributes: (i) nucleotide identity (A, T, C, G), (ii) codon phase (1st, 2nd, or 3rd position), (iii) encoded amino acid, and (iv) synonymous codon identity.

To quantitatively validate these observations, we analyzed an independent data set of 3,765 viral sequences from held-out development and test sets. We employed permutational multivariate analysis of variance (PERMANOVA) (*63*) to test for significant differences in embedding clusters corresponding to nucleotide identity, codon position, and coding semantics. Furthermore, we conducted linear probing experiments to assess the linear separability of these biological features, in which a logistic regression classifier was trained on the frozen embeddings. We benchmarked LucaVirus against state-of-the-art genomic models, including Nucleotide Transformer (*30*), Evo (*14*), Evo2 (*13*), and LucaOne (*20*).

Finally, we evaluated whether LucaVirus captures virus-enriched genomic features associated with programmed ribosomal frameshifting (PRF). Specifically, 300-bp windows centered on annotated PRF sites were extracted from the FSDB database (*31*). We evaluated whether embedding’s ability to distinguish the relative position of nucleotides within a codon (codon phase). A correctly learned representation should reflect the phase shift inherent in the biological mechanism; failure to identify the frameshift would result in misaligned phase predictions. We compared the codon phase discrimination performance of LucaVirus against baseline models using the linear probing method described above.

**Protein Representation and Family Discrimination.** To evaluate protein-level representation capacity, we compared LucaVirus against mainstream models trained across all domains of life (including eukaryotic viruses, which are underrepresented or partially excluded in some models), namely, LucaOne (*20*), ESM2 (*11*), and ESMC (*12*). Viral reference protein sequences were retrieved from NCBI RefSeq and grouped into families using the VOGDB pipeline (*64*), which employs DIAMOND (*38*) for pairwise alignment followed by Markov clustering (*65*). We selected clusters containing >20 sequences and randomly sampled 10 families, from which 10 sequences were randomly sampled (N=100 total), and per-residue feature embeddings were extracted. PCA was then applied to visualize separation of protein families within the learned representation space.

**Correlation with Evolutionary Divergence.** To quantify the relationship between LucaVirus embeddings and evolutionary divergence, we performed correlation analyses between embedding space distances (cosine and Euclidean) and genetic divergence metrics (p-distance and maximum likelihood estimates of substitution rates). The primary data set comprised 5,566 viral orthologous protein families with corresponding coding DNA sequences (CDS) from RefSeq. To evaluate generalization to previously “unseen” viral diversity, we additionally included 938 protein families from novel RNA viruses (“dark matter” viruses) identified in our prior study (*10*), which exhibit no significant BLAST hits to the pre-training corpus. Protein families were defined using *de novo* clustering via the VOGDB pipeline (*64*). For each family, mean pairwise embedding distances were computed and correlated with genetic distances derived from multiple sequence alignments (MSA).

**LMAlign: Embedding-Based Sequence Alignment and Validation.** To assess whether LucaVirus representations capture deep evolutionary signals, we developed LMAlign, an embedding-based pairwise sequence alignment algorithm. Input sequences are first processed by LucaVirus to generate dense embedding matrices of size 𝐿 × 2560. A dynamic similarity matrix (𝐿_1_ × 𝐿_2_) is then computed using cosine similarity between residue-level embeddings, which serves as the scoring matrix for dynamic programming. This framework enables optimal local and global alignments analogous to the Smith-Waterman and Needleman-Wunsch algorithm respectively, but directly from the model’s semantic feature space.

LMAlign was benchmarked for remote homology detection using the SCOPe 2.08 data set (*33*) filtered for viral proteins. To evaluate performance across varying degrees of evolutionary distance, test pairs were stratified across three hierarchical levels: family, superfamily, and fold. We performed all-against-all pairwise alignments, comparing LMAlign (local and global modes) with conventional Needleman-Wunsch and Smith-Waterman alignments utilizing BLOSUM62 (*32*) and BLASTP (*66*) scores. Performance was evaluated using receiver operating characteristic (ROC) curves and errors per query (EPQ) metrics, quantifying sensitivity in detecting homologous relationships beyond the reach of sequence-identity-based methods.

### A General Modeling Framework for Downstream Tasks

**Unified Model Architecture.** We implemented a standardized framework to adapt pre-trained models for diverse downstream applications. The core architecture consists of a frozen pre-trained encoder (e.g., LucaVirus) for embedding extraction, followed by a trainable value-level attention pooling (VALP) layer (*67*) and a fully connected (FC) prediction head. This design concentrates task-specific learning in a lightweight head while preserving the generality and stability of the pre-trained backbone. The framework supports two input modes: (1) Single-Sequence Mode: Processes individual sequences (e.g., for function prediction). It combines the frozen embedding pathway with an optional, trainable lightweight Transformer encoder to capture task-specific features. (2) Multi-Sequence Mode: Processes interacting sequences (e.g., for antibody-antigen binding). Each sequence is encoded and pooled independently; the resulting vectors are concatenated to form a joint representation. The final FC head is flexible and supports binary, multi-class, or multi-label classification, as well as regression by adjusting output dimensions.

**Training and Reporting Protocol.** To ensure rigorous and unbiased evaluation, all data sets were strictly partitioned into independent training, development, and test sets. The model was trained using the AdamW optimizer. For each task, we conducted a grid search over learning rates (1×10^-4^, 2×10^-4^) and batch sizes (8, 16) to determine optimal training hyperparameters. Model selection was based on optimal performance on the development set, and final results are reported on the independent test set.

### Challenge 1: Identification of Viral “Dark Matter”

**Data Set and Task Setup.** We formulated two binary classification tasks targeting viral hallmark proteins: RNA-dependent RNA polymerase (RdRp) and Capsids. The data set comprises 5,979 RdRp positives and 229,434 negatives sourced from previous studies, alongside 216,742 Capsid positives (UniProt KW2013) balanced against 83,461 hard negatives (e-value < 10^-5^ via DIAMOND (*38*) alignment against UniProtKB (*60*) and NCBI nr databases) and 133,281 easy negatives randomly sampled from non-capsid viral proteins on SwissProt. The data is randomly split into 75% training and 10% development, 15% testing sets. We employed the general modeling framework for downstream tasks described above configured with a binary classification head for these tasks.

**Validation on Viral “Dark Matter”.** To assess the model’s capacity to detect remote viral homologs effectively (“dark matter”), we screened sequences from the LucaProt data set (*10*) that lacked significant matches in the NCBI nr database. Candidates predicted as positive were clustered using DIAMOND BLASTP (*38*) and MCL (*65*). Representative sequences from each cluster were subjected to structure prediction using ESMFold (*11*) and ColabFold (*41*), followed by FoldSeek (*37*) searches against PDB (*68*), UniRef50, and SwissProt (*60*) (e-value < 1e-5) to verify structural homology to known capsids. High-confidence candidates were further validated via high-accuracy structure prediction using AlphaFold3 (*48*).

**Benchmarking against Bioinformatic Approaches.** To rigorously evaluate LucaVirus against standard bioinformatics pipelines, we conducted a comparative benchmark using sequence similarity and structure-based search tools. To eliminate data leakage and test generalization, we strictly removed all test sequences sharing >50% sequence identity with the pre-training corpus. We compared LucaVirus against DIAMOND BLASTP (ultra-sensitive mode) (*38*), MMseqs2 profile search (sensitivity −s 7) (*40*), HHblits (10 iterations) (*39*), and FoldSeek (using ProstT5 module (*69*) for structure prediction) (*37*). To ensure a fair comparison, the database for all retrieval methods was constructed exclusively from the LucaVirus training set. For the above methods, we defined two classification criteria using an e-value threshold 10^-5^: (1) Top-hit Mode: Classification is determined solely by the hit with the highest bitscore. If the top hit is a capsid, the query is classified as positive. (2) Strict Consensus Mode: A query is classified as positive only if all significant hits (e-value < 10^-5^) are annotated as capsids; otherwise, it is classified as negative.

**Computational Efficiency Analysis.** To evaluate scalability and runtime performance, we recorded the mean per-sequence query time and total database preparation/training time. To ensure a fair comparison between AI-driven and traditional algorithms, we utilized cost-equivalent hardware configurations: the AI models were tested on a single NVIDIA A100 (80GB) GPU, while CPU-based bioinformatics tools utilized dual-socket AMD EPYC 7H12 CPUs (128 cores total), with all tools configured to use 128 threads to fully saturate the CPU resources.

### Challenge 2: Characterization of Unknown Proteins

**Data Set and Task Setup.** We formulated a multi-label classification task utilizing 10,715 viral proteins from UniProtKB (*60*) annotated with Enzyme Commission (EC) numbers, filtering for labels with at least 100 associated sequences to ensure enough training samples per label. 70% training and 15% development, 15% testing sets. The model employed the general modeling framework for downstream tasks, configured with a sigmoid activation multi-label prediction head to output probabilities for enzyme classes.

**Benchmarking on Structure-Inferred Annotations.** To evaluate the model’s capacity to annotate proteins that lack sequence homology to known enzymes, we utilized the structurally defined viral data set from ref (*8*). In this data set, ground truth labels were generated by propagating InterProScan (*70*) functional annotations (CDD (*71*), TIGRFAM (*72*), Pfam (*73*)) within structural clusters defined by ColabFold (*41*) and FoldSeek (*37*) (propagation threshold: >25% intra-cluster consistency). To ensure a rigorous test of generalization, we strictly excluded all test sequences sharing >50% sequence identity with the LucaVirus pre-training corpus. We compared LucaVirus against ESM2-3B (*11*) and the original structure-based propagation method, reporting standard classification metrics (accuracy, precision, recall, F1) as well as annotation coverage—defined as the proportion of total proteins successfully assigned a specific EC label.

**Experimental Validation of Putative Proteases.** To validate the model’s predictions, RNA viral proteins predicted to be proteases (EC 3.4.-.-) from the LucaProt study, which identified distant homologs of dark matter RNA virus proteins, were selected. Candidates were clustered into families (following the VOGDB methodology (*64*)), and two representatives from distinct families: CL6_PICL_PRO (22.4 kDa) and CL31_SOBL_PRO (21.7 kDa), were selected for characterization.

**Protein Expression and Purification.** Genes were cloned into expression vectors and transformed into *Escherichia coli* BL21 (DE3). Cultures were grown in LB medium at 37°C until OD_600_ ≈ 0.6, induced with 0.5 mM IPTG, and incubated at 16°C for 16 hours. Cells were harvested and resuspended in Lysis Buffer (20 mM Tris-HCl, 400 mM NaCl, 20 mM imidazole, pH 7.5). Following high-pressure homogenization (700 Pa) and clarification by centrifugation (16,000 rpm, 25 min, 4°C), the supernatant was loaded onto nickel affinity resin. After washing with 10 column volumes of Lysis Buffer, the 12His-SUMO tag was cleaved on-column by overnight protease digestion in Elution Buffer (50 mM Tris-HCl, 100 mM NaCl, pH 7.0) to elute the native protein.

**Functional Assays.** Protein purity was verified via SDS-PAGE on 4–20% pre-cast gels. Proteolytic activity was quantified using a fluorescence-based protease assay (Beyotime, Catalog No. P0403S) according to the manufacturer’s protocol.

### Challenge 3: Forecasting Virus Evolvability

**Supervised Fitness Prediction.** We formulated the prediction of viral fitness landscapes as a scalar regression problem utilizing the general modeling framework for downstream tasks. To rigorously evaluate the model’s capacity to forecast evolutionary trajectories across distinct viral families and selection pressures, we curated three comprehensive Deep Mutational Scanning (DMS) data sets:

**SARS-CoV-2 Receptor Binding (Temporal Generalization).** We utilized DMS data measuring the ACE2 binding affinity of the SARS-CoV-2 receptor-binding domain (RBD) (*74*). To simulate a realistic forecasting scenario, we adopted a temporal split strategy: the model was trained on early variants (Wuhan-Hu-1, Alpha, Beta; N=34,055) and evaluated on later-emerging variants (Delta, Iota; N=19,274). This design tests the ability to generalize fitness predictions to future evolutionary lineages.

**Influenza H3N2 Immune Escape (Antigenic Generalization).** To assess predictions of antigenic evolution, we incorporated the immune escape landscape of Influenza A virus (H3N2, strain A/Hong Kong/19/2014) (*75*). The fitness metric is defined as the **e**scape fraction, calculated as the ratio of mutant abundance in serum-selected cultures versus non-selected controls. We implemented a cross-serum split: the model was trained on escape profiles derived from specific sera and tested on differing sera to evaluate its ability to capture transferable antigenic features independent of specific antibody repertoires.

**Enterovirus A Polyprotein (Indel Robustness).** To extend prediction capabilities beyond substitutions, we utilized a Deep Indel Scanning data set for the Enterovirus A polyprotein (*76*). This data set measures fitness via relative abundance in cell culture following the introduction of single amino acid substitutions, insertions, and deletions. Given the high dimensionality and lack of replicates for specific mutants, we employed a randomized split (75% training, 10% development, 25% testing) to assess the model’s robustness to structural perturbations including indels.

**Zero-Shot Fitness Estimation.** To benchmark the model’s intrinsic, unsupervised understanding of viral fitness, we performed zero-shot predictions on the viral subset of the ProteinGym benchmark (*44*), comprising 31 diverse viral protein assays. Consistent with established methodologies, fitness scores were computed as the masked token Log-Likelihood Ratio (LLR) between the mutant and wild-type sequences:

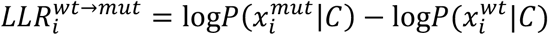

Where 𝑖 denote the 𝑖^th^ amino acid within protein sequence, 𝑤𝑡 denote wildtype residue, 𝑚𝑢𝑡 denote mutant residue, 𝑃(𝑥|𝐶) represents the model’s predicted probability of masked token 𝑥 given the unmasked context 𝐶.

### Challenge 4: Prediction of Antigen-Antibody Binding Affinity

**Data Set and Task Setup.** We constructed a binary classification data set using positive samples from CoVAbDab (*47*) (human-derived, strong SARS-CoV-2 binders) and negative samples augmented from pre-pandemic human BCR sequences (assumed non-binders, 10x positive count). The data was split into 70% training and 30% testing sets. We employed the general modeling framework described above, concatenating pooled embeddings from antibody heavy chain, light chain, and antigen sequences to predict binding probability.

**Real-World Experimental Validation.** We established a validation pipeline from single-cell sequencing to functional ELISA. Single-cell V(D)J data from convalescent/vaccinated donors was processed (Cell Ranger, IgBLASTn (*77*)) to identify paired heavy/light chains. To prevent data leakage, we strictly excluded candidates with <10 amino acid mismatches relative to the training set. Selected candidates were synthesized, expressed in HEK293T cells, and purified using Protein-A agarose beads. Binding affinity was assessed via ELISA assay. High-binding microtiter plates (Costar) were coated with 2 µg/mL of SARS-CoV-2 WT Spike, WT RBD, or Omicron Spike recombinant proteins overnight at 4°C. Following blocking (3% BSA), purified mAbs (10 µg/ml, 1:3 serial dilution) were incubated for 1 hour at 37°C. Bound antibodies were detected using HRP-conjugated goat anti-human IgG (1:2000). OD_405_ exceeding twice the PBS blank control was defined as positive binding. Binding strength (AUC) was visualized via heatmaps generated in GraphPad Prism 8.0.

We utilized this in-house data set to benchmark LucaVirus against generalist protein language models (e.g., ESM2 (*11*), ESMC (*12*)). Additionally, we evaluated two quantitative structure-based prediction approaches: (1) AlphaFold 3+CSM-AB, a composite pipeline combining AlphaFold 3 (*48*) for structural modeling and docking with the CSM-AB (*49*) scoring function; and (2) DG-Affinity (*50*). For these methods, we compared the predicted binding energies (or affinity scores) for experimentally validated binders versus non-binders. Statistical significance of the discrimination capability was assessed using independent *t*-tests.

**Comparative Benchmarking against Specialized Models.** We further benchmarked LucaVirus against three distinct classes of state-of-the-art models: (1) mBLM (protein language model) (*51*): Evaluated on our in-house experimental data set. We aggregated mBLM’s multi-class outputs into a binary binding prediction to facilitate direct comparison. (2) MAGE (generative language model) (*53*): Evaluated on 20 *de novo* antibodies generated by MAGE, predicting their binding to Wuhan-Hu-1 RBD/Spike. To assess the relative difficulty of this task, we confirmed that the sequence identity of these test antibodies to the MAGE training set was higher than to the LucaVirus training set, implying a stricter test of generalization for LucaVirus. (3) AF2Complex (structure-based) (*52, 78*): Evaluated on 971 antibodies (471 binders, 500 non-binders) from the AF2Complex study, filtered to exclude sequences with <10 mismatches relative to our training data. We computed optimal thresholds (Youden’s index) for AF2Complex’s continuous interaction scores to enable binary classification metric comparison.

## Acknowledgements

We thank J.-S. Eden for helpful discussions and assistance with manuscript preparation. The computations in this research were supported by the CFFF platform of Fudan University and the High-performance Computing Platform of YZBSTCACC (YaZhou Bay Science and Technology City Advanced Computing Center).

## Author contributions

Conceptualization: Y-FP, YH, Z-RL, MS; Methodology: Y-FP, YH; Investigation: Y-FP, YH, Y-QL, S-NL; Project administration: BL, Y-QC, Z-RL, MS; Supervision: BL, Y-QC, Z-RL, MS; Writing – original draft: Y-FP; Writing – review & editing: All authors.

## Data and materials availability

All data, code, and materials used in this analysis are publicly available to researchers for reproducing or extending the findings. The source code for modeling and pretraining, along with instructions for obtaining pretrained weights, is accessible at https://github.com/LucaOne/LucaVirus. The code for downstream task modeling and instructions for retrieving trained model weights are available at https://github.com/LucaOne/LucaVirusTasks. Pretrained weights for the LucaVirus foundation model, downstream models, and the training data set are accessible at http://47.93.21.181/lucavirus/ and on Zenodo (DOI: 10.5281/zenodo.15703216).

