## Supplementary Materials for "Predicting the Evolutionary and Functional Landscapes of Viruses with a Unified Nucleotide-Protein Language Model: LucaVirus"

**The PDF file includes:**

Figs. S1 to S11
Tables S1 to S13

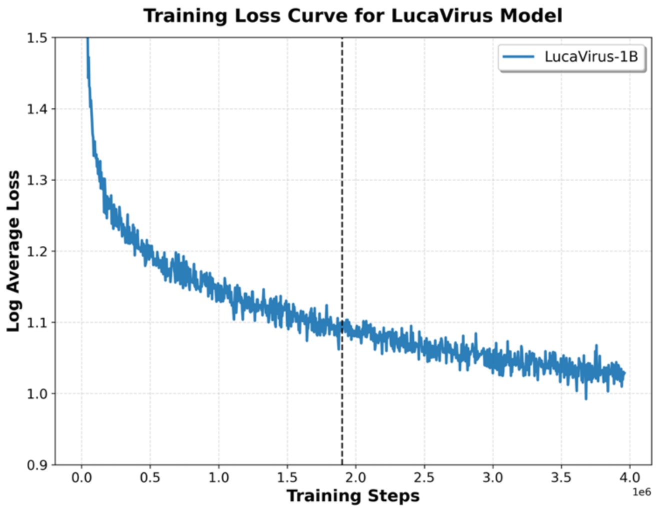


**Fig. S1. Pretraining loss curve for LucaVirus-1B model.** We trained LucaVirus using eight A100 GPUs on Alibaba Cloud for a total of 3,800,000 steps (approximately 1.9 epochs) over a duration of 70 days. 'Log Average Loss' refers to the average loss calculated every 4,000 steps. Dashed line indicates the end of first epoch.

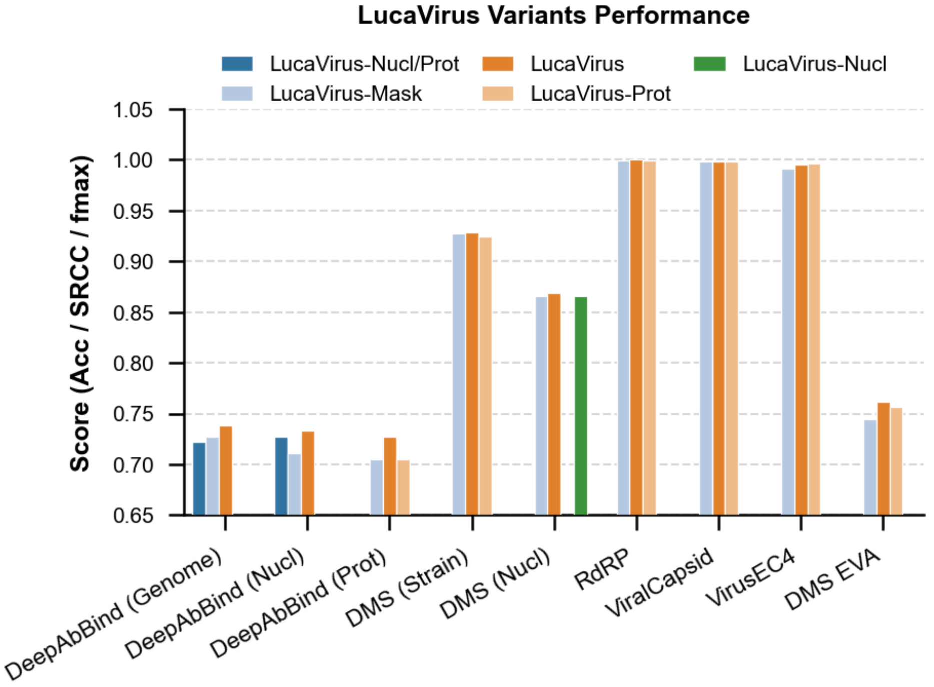


**Fig. S2. Ablation analysis of the LucaVirus architecture.** The impact of individual components is assessed across task types using standard metrics: accuracy for classification, Fmax for multi-label classification, and Spearman's correlation coefficient (SRCC) for regression. Results identify critical elements for model efficacy. Refer to Table S3 for detailed data. Note: The antibody binding performance metrics in this table are based on all 98 antibodies, including those with weak binding affinity.

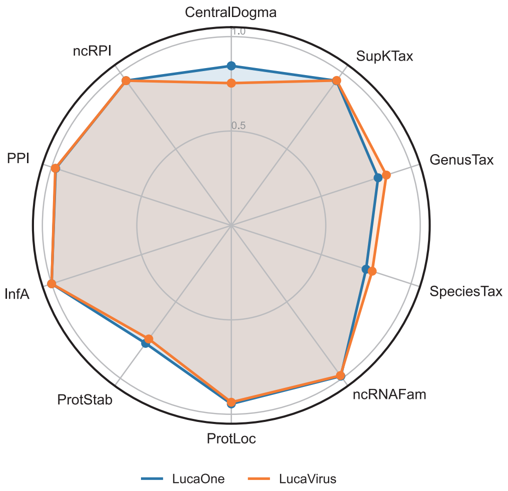


**Fig. S3. Generalization assessment of LucaVirus on a multi-modal benchmark of biological sequence tasks.** LucaVirus, trained on viral sequences, is compared to LucaOne on 10 original benchmark tasks spanning protein and nucleotide prediction. Performance metrics are classification accuracy and Spearman's correlation for regression. The metrics for LucaOne were obtained at the 5,600,000-step checkpoint from LucaOne (20). The comparison underscores the model's broad applicability. Refer to Table S4 for a complete task breakdown.

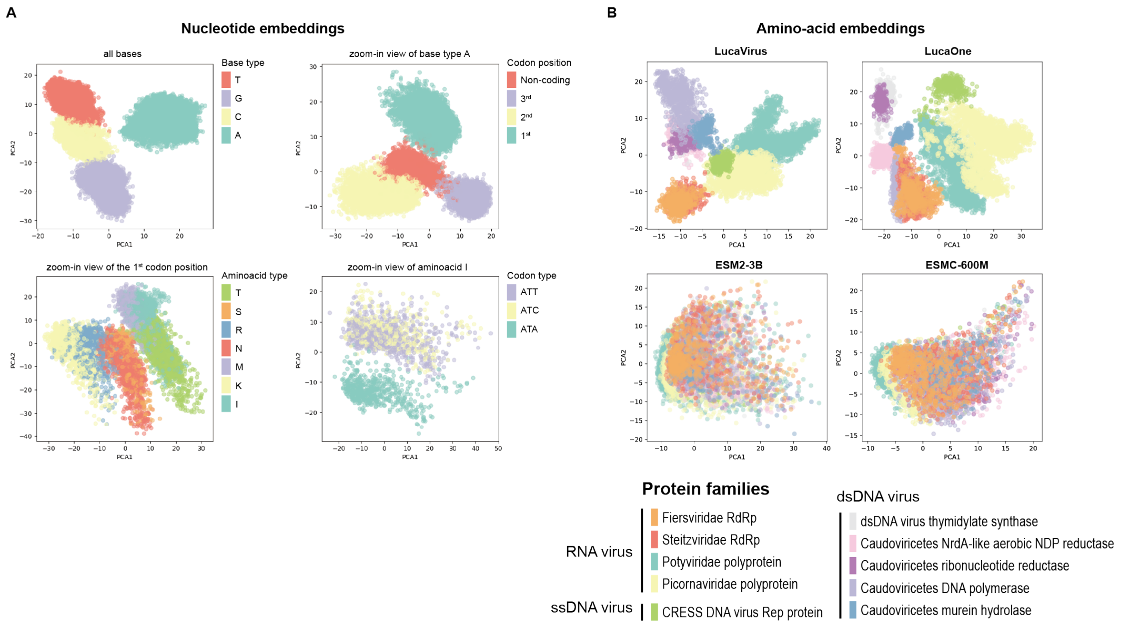


**Fig. S4. LucaVirus captures the relationship between nucleotide and protein sequences at single amino acid resolution.** (**A**) Nucleotide sequence features learned by LucaVirus. This panel visualizes the embedding matrix from 30 enterovirus reference genomes processed by LucaVirus, reduced via PCA. Each point represents a nucleotide token's embedding vector. The four sub-plots are: (i) PCA of all nucleotide tokens, colored by base type. (ii) PCA of adenine (A) tokens, colored by codon position. (iii) PCA of adenine tokens at the first codon position, colored by encoded amino acid. (iv) PCA of adenine tokens at the first position of codons encoding isoleucine (I), colored by codon type. (**B**) Viral protein feature spaces from different language models. This panel compares how LucaVirus, LucaOne, ESM2-3B, and ESMC-600M represent viral protein sequences from 100 proteins across 10 families. Embeddings are reduced via PCA, with points colored by protein family. Each point represents an amino acid token's embedding vector.

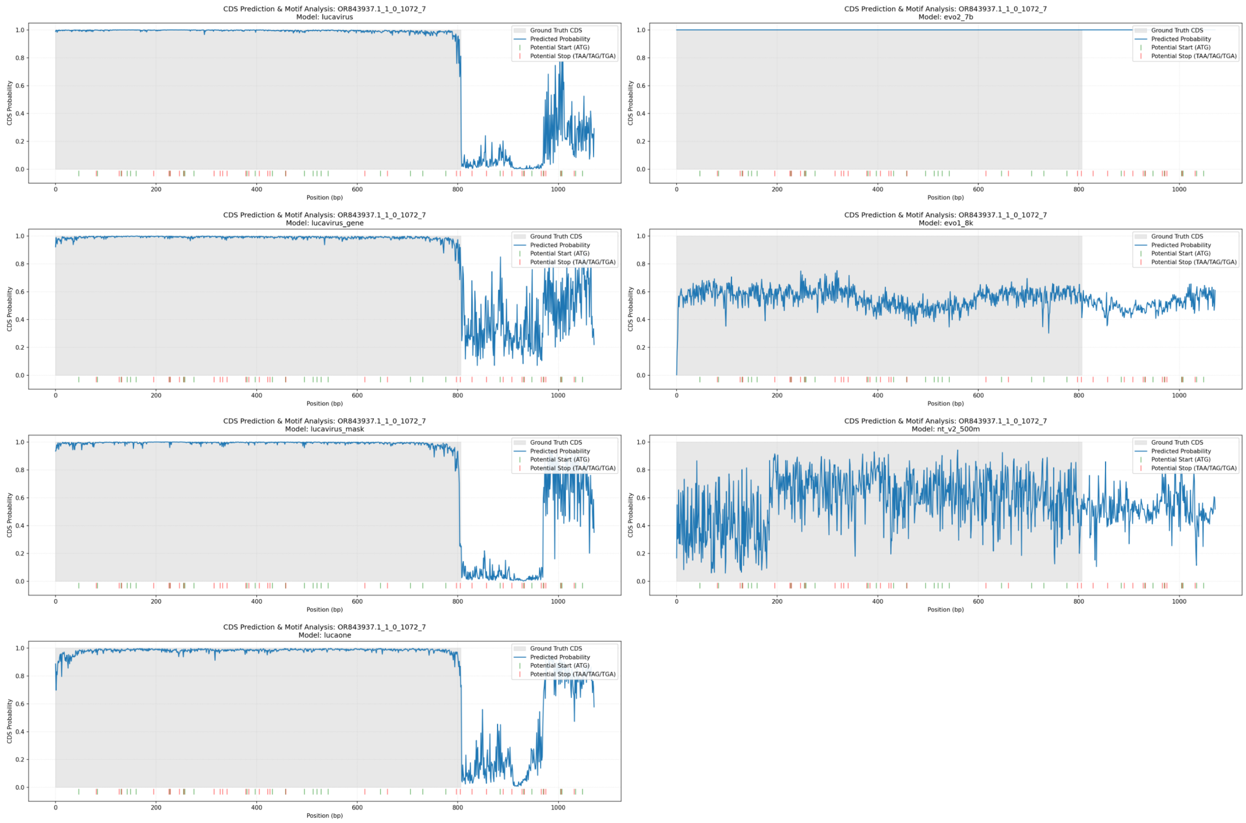


**Fig. S5. Comparative visualization of linear probe predictions for coding sequence (CDS) identification.** The plot depicts model outputs for the held-out exemplar OR843937.1. Gray shading denotes the ground-truth CDS region per NCBI annotation. The green and red line indicate location of start and stop codon respectively. The continuous blue trace for each model (LucaVirus, LucaVirus-Nucl, LucaVirus-Mask, LucaOne, Evo2, Evo1, Nucleotide Transformer) represents the per-position probability of being within a CDS, with a classification threshold at 0.5.

The results demonstrate that the Luca-family models successfully identified a non-complete coding region (clipped at the 5’-end, lacking a canonical start codon) while correctly ignoring other non-coding open reading frames (ORFs). Among these, LucaVirus achieved the highest accuracy and cleanest signal-to-noise ratio. In contrast, the Evo-family and Nucleotide Transformer models failed to accurately resolve the CDS boundaries or distinguish between coding and non-coding regions.

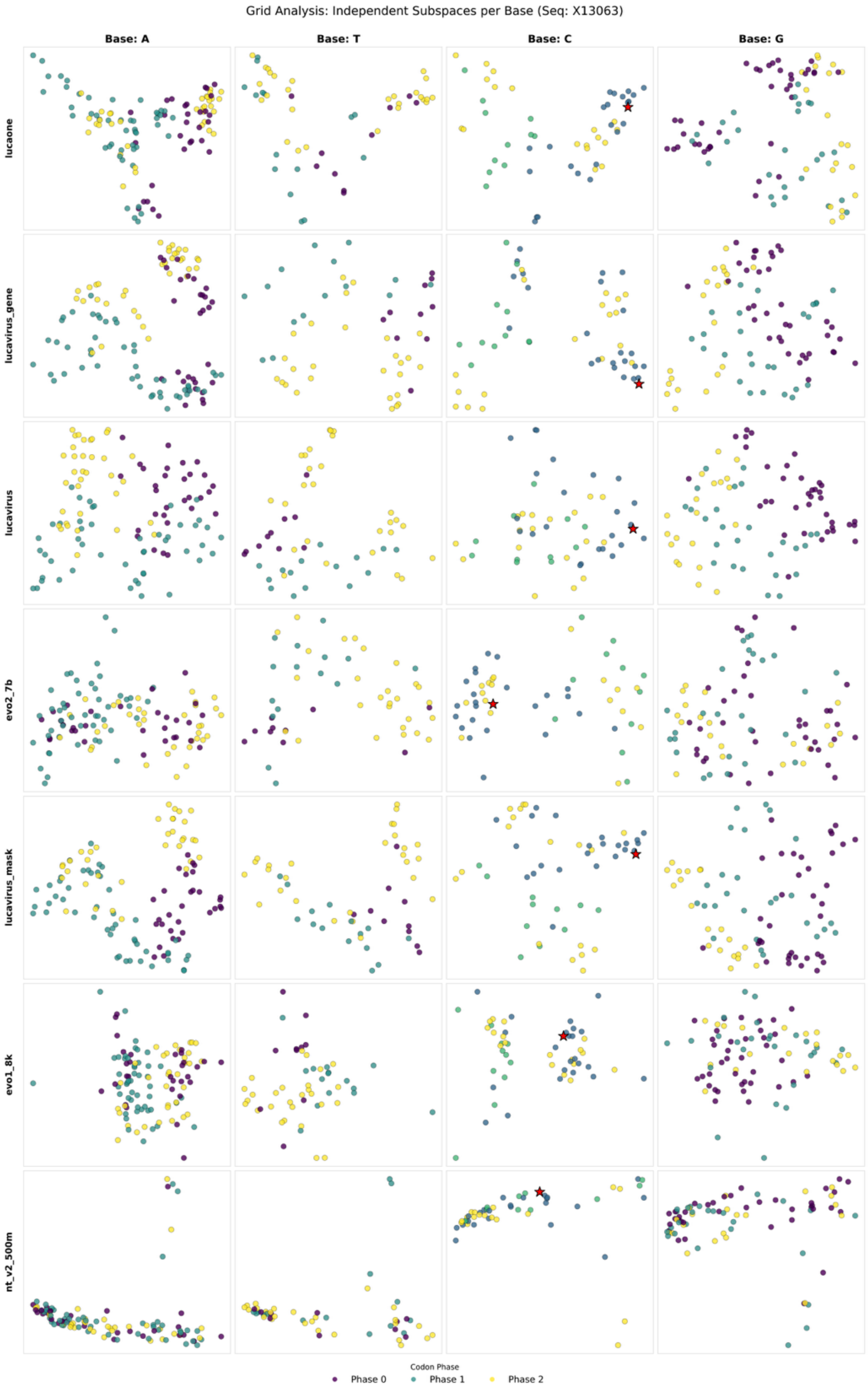


**Fig. S6. Comparative visualization of LLM embeddings near a programmed ribosomal frameshifting (PRF) site.** This grid analysis illustrates the embedding space for sequence X13063, specifically focusing on the region surrounding a -1 PRF event. The columns represent independent subspaces for each nucleotide base (A, T, C, G), while rows display results for different models: LucaOne, LucaVirus-Nucl, LucaVirus, Evo2-7b, LucaVirus-Mask, Evo1-8k, and NT-v2-500m.

Points are colored by codon phase (Phase 0: purple; Phase 1: teal; Phase 2: yellow), which represents the relative position of a base within a triplet codon. The red star indicates the programmed ribosomal frameshifting (slippery) site, where the ribosome shifts the reading frame by one base, thereby altering the subsequent codon phases.

The results demonstrate that the LucaVirus and related Luca-family embeddings effectively capture the local phase structure, shown by the distinct clustering of phases even across the frameshifting transition. In contrast, models like Evo2-7b, Evo1-8k, and NT-v2-500m exhibit significant mixing of phases in their embedding spaces, suggesting a failure to maintain a clear representation of the translational reading frame in the vicinity of non-canonical translation events.

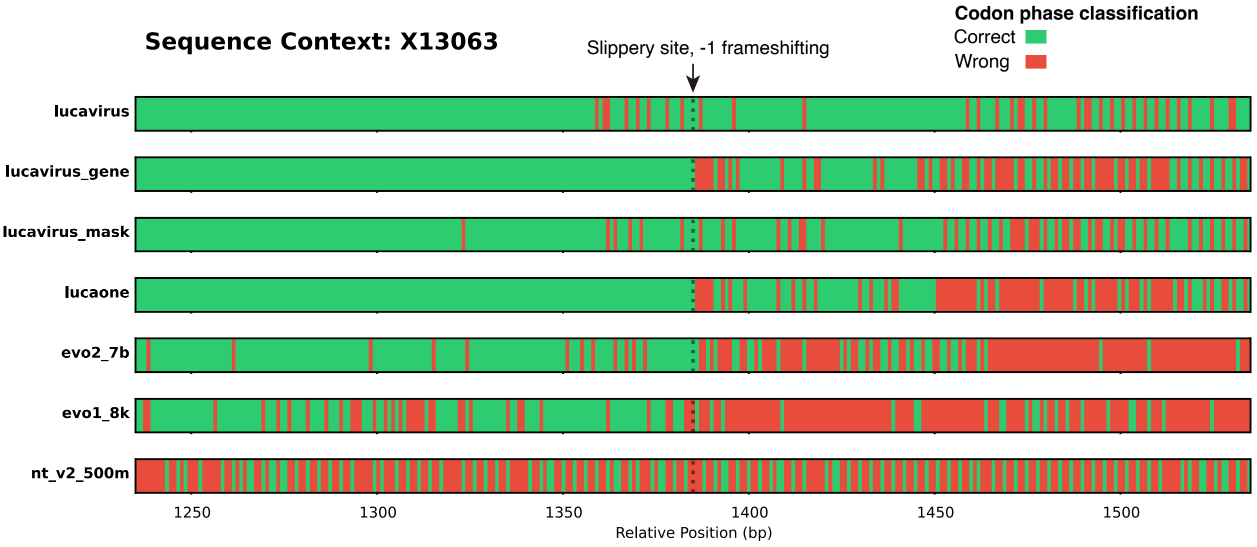


**Fig. S7. Codon phase classification accuracy across a programmed ribosomal frameshifting (PRF) site.** Complementing the embedding analysis in Fig. S6, this figure displays the linear probe prediction results for sequence X13063. The vertical dashed line and arrow pinpoint the slippery site (at approximately 1380 bp) where a -1 PRF event occurs. Each horizontal track represents the performance of a specific model, with green segments indicating correct codon phase identification and red segments indicating misclassification.

The results reveal that LucaVirus and related Luca-family models maintain high classification accuracy both before and after the frameshifting event, demonstrating that their embeddings successfully capture the transition in the translational reading frame. In contrast, Evo2-7b, Evo1-8k, and NT-v2-500m exhibit a marked increase in errors (dense red regions) following the slippery site, suggesting that these models fail to adapt to non-canonical translational shifts. These findings highlight the superior capability of the Luca-family models in representing complex viral genomic features.

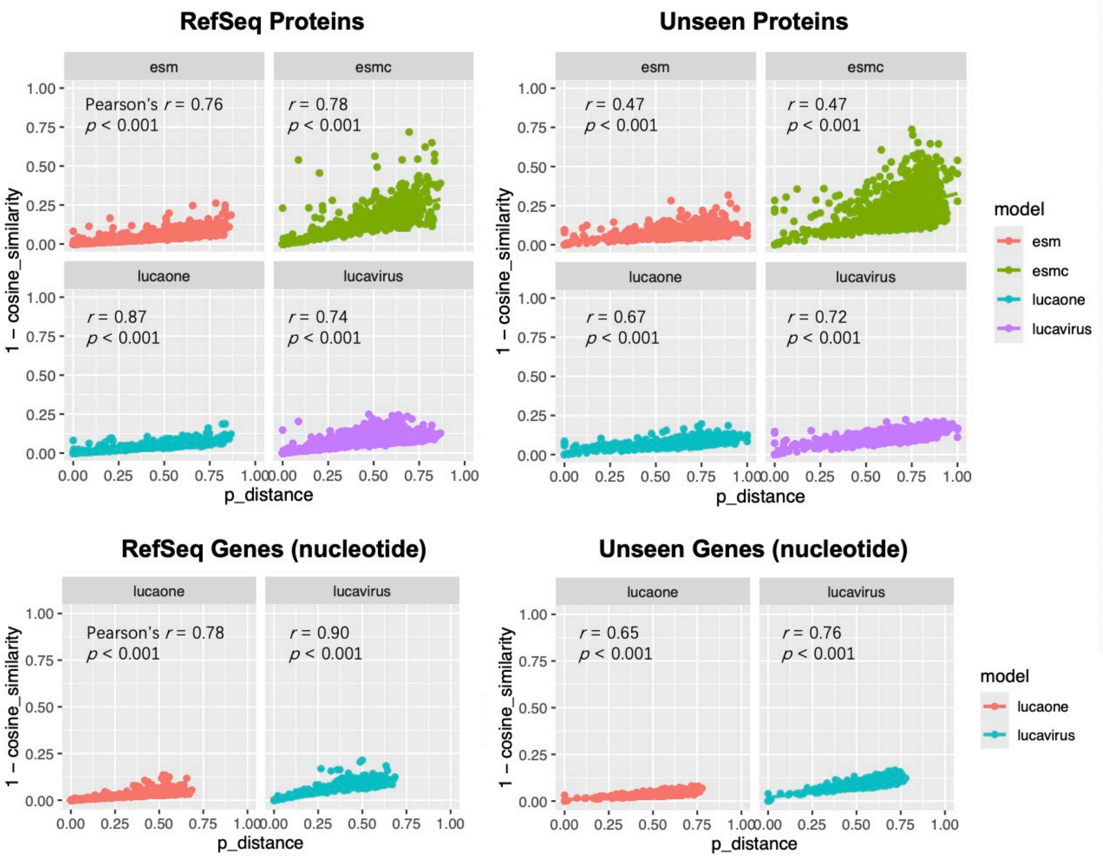


**Fig. S8. Correlation between the mean cosine distance of sequence embeddings and p-distance (1 – percent identity).** Among the popular LLMs tested, LucaVirus demonstrates the most robust and generalizable relationship with sequence divergence, maintaining strong performance even on previously unseen protein data sets. By contrast, the other three models show a marked drop in performance outside of their training data.

**Algorithm 1** Embedding-based Smith-Waterman with Affine Gaps

**function** EMB_SW_ALIGN_AFFINE$(S, T, \sigma, \mu, \alpha, \beta)$
    **Input:** Similarity matrix $S$ $(m \times n)$, threshold $T$, optional match score $\sigma$, mismatch penalty $\mu$, gap open penalty $\alpha$, gap extend penalty $\beta$
    **Output:** Aligned indices, alignment score
    **Construct scoring matrix $W$:**
    **for** each $i, j$ **do**
        **if** $S_{i,j} > T$ **then**
            **if** $\sigma =$ None **then**
                $W_{i,j} \leftarrow S_{i,j}$
            **else**
                $W_{i,j} \leftarrow \sigma$
            **end if**
        **else**
            $W_{i,j} \leftarrow \mu$
        **end if**
    **end for**
    Initialize $M, X, Y, H$ as $(m+1) \times (n+1)$ zero matrices
    max_score $\leftarrow 0$, max_pos $\leftarrow (0, 0)$
    **for** $i = 1$ to $m$ **do**
        **for** $j = 1$ to $n$ **do**
            $M_{i,j} \leftarrow \max(0, M_{i-1,j-1} + W_{i,j}, X_{i-1,j-1} + W_{i,j}, Y_{i-1,j-1} + W_{i,j})$
            $X_{i,j} \leftarrow \max(0, M_{i-1,j} + \alpha + \beta, X_{i-1,j} + \beta)$
            $Y_{i,j} \leftarrow \max(0, M_{i,j-1} + \alpha + \beta, Y_{i,j-1} + \beta)$
            $H_{i,j} \leftarrow \max(M_{i,j}, X_{i,j}, Y_{i,j})$
            **if** $H_{i,j} >$ max_score **then**
                max_score $\leftarrow H_{i,j}$
                max_pos $\leftarrow (i, j)$
            **end if**
        **end for**
    **end for**
    Perform standard Smith-Waterman traceback with affine gaps from max_pos
    **Return** the alignment and max_score
**end function**

**Fig. S9. The pseudo-code for the LMAlign algorithm – an embedding-based Smith-Waterman alignment method.**

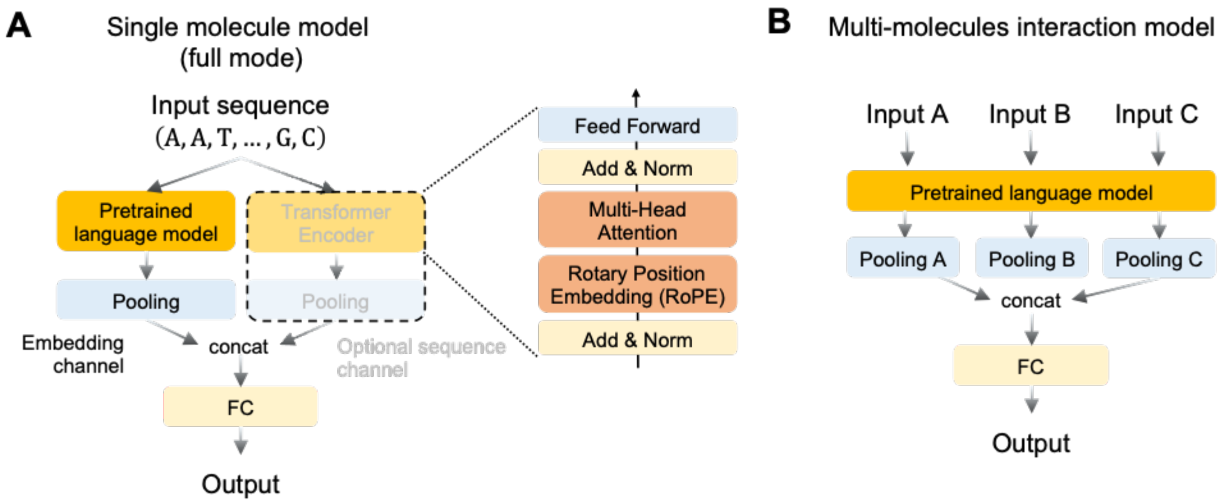


**Fig. S10. General framework for using pretrained language models for specific downstream tasks.** In all tasks examined in this study, the weights of the pretrained language model were kept frozen during downstream training.

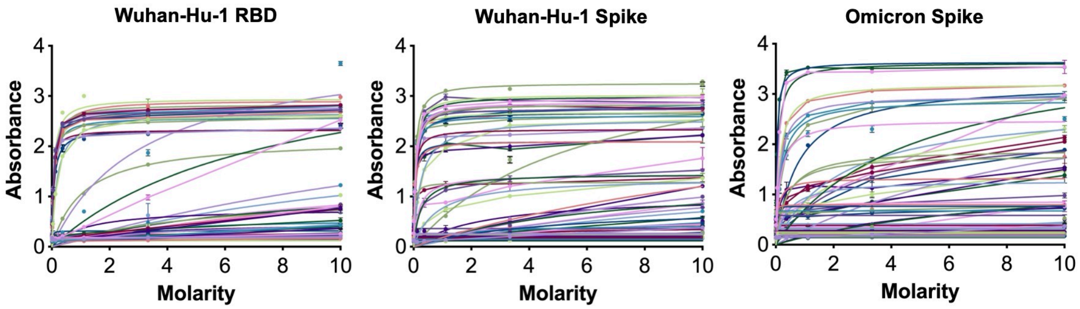


**Fig. S11. Validation of antibody-antigen binding using ELISA.** We randomly selected 98 antibodies obtained through single B cell sequencing from convalescent COVID-19 patients in our previous study. ELISA assays were performed to assess the binding of these antibodies to the receptor-binding domain (RBD) and full-length Spike protein of the Wuhan-Hu-1 strain, as well as the full-length Spike protein of the Omicron variant.

**Table S1. Composition of pre-training data set**

| Source | Type | Token counts | Sequence counts | Seq-level annotation | Span-level annotation |
|---|---|---|---|---|---|
| NCBI Virus | Nucleotide | 2,322,433,600 | 3,233,945 | taxonomy | CDS |
| Wolf et al. (ref *22*) | Nucleotide | 1,121,406,280 | 1,875,616 | n/a | n/a |
| Gregory et al. (ref *24*) | Nucleotide | 1,465,432,141 | 1,875,225 | n/a | n/a |
| Hou et al. (ref 9) | Nucleotide | 891,253,024 | 373,079 | n/a | n/a |
| GPD (ref *21*) | Nucleotide | 349,588,480 | 2,910,001 | n/a | n/a |
| Neri et al. (ref *20*) | Nucleotide | 393,776,802 | 168,727 | n/a | n/a |
| Zayed et al. (ref *23*) | Nucleotide | 28,931,582 | 29,112 | n/a | n/a |
| UniProt-Trembl | Protein | 1,627,140,177 | 5,164,087 | taxonomy/keywords | Super family/domain/site |
| UniProt-SwissProt | Protein | 7,779,199 | 17,212 | taxonomy/keywords | Super family/domain/site |
| UniProt-UniRef50 | Protein | 141,972,000 | 605,379 | taxonomy/keywords | Super family/domain/site |
| ColabFold-envdb | Protein | 75,973,285 | 282,640 | taxonomy | n/a |

**Table S2. Model configuration of six LucaVirus pre-trained variants.**

| Pretrained model | Parameters | Embedding dimension | Pretraining tasks | Modality |
|---|---|---|---|---|
| LucaVirus-Large | 1B | 2560 | MLM + semi-supervised tasks | Protein/Nucleotide |
| LucaVirus-Base | 112M | 256 | MLM + semi-supervised tasks | Protein/Nucleotide |
| LucaVirus-Small | 62M | 128 | MLM + semi-supervised tasks | Protein/Nucleotide |
| LucaVirus-Mask | 1B | 2560 | MLM only | Protein/Nucleotide |
| LucaVirus-Prot | 1B | 2560 | MLM + semi-supervised tasks | Protein |
| LucaVirus-Nucl | 1B | 2560 | MLM + semi-supervised tasks | Nucleotide |

**Table S3. Detailed performance metrics for model ablation studies.**

| Task | Model | Classification metrics | | | | | Multi-label metrics | | Regression metrics | |
|---|---|---|---|---|---|---|---|---|---|---|
| | | Acc | F1 | AUC | PR-AUC | MCC | Jaccard | Fmax | Spearman | Pearson |
| DeepAbBind v2 (genome) | LucaVirus-Nucl/Prot | 0.7222 | 0.5192 | **0.6950** | 0.5801 | 0.3464 | | | | |
| | LucaVirus-Mask | 0.7277 | 0.5242 | 0.6648 | **0.5862** | 0.3589 | | | | |
| | LucaVirus | **0.7388** | **0.5765** | 0.6913 | 0.5529 | **0.3972** | | | | |
| DeepAbBind v2 (nucl) | LucaVirus-Nucl/Prot | 0.7277 | 0.5148 | 0.6800 | 0.5801 | 0.3566 | | | | |
| | LucaVirus-Mask | 0.7111 | 0.5357 | 0.6319 | 0.5113 | 0.3335 | | | | |
| | LucaVirus | **0.7333** | **0.5385** | **0.6951** | **0.6024** | **0.3741** | | | | |
| DeepAbBind v2 (original) | LucaVirus-Prot | 0.7055 | 0.5046 | 0.6755 | 0.5918 | 0.3104 | | | | |
| | LucaVirus-Mask | 0.7055 | 0.4952 | 0.6603 | 0.5179 | 0.3067 | | | | |
| | LucaVirus | **0.7278** | **0.5812** | **0.7124** | **0.6107** | **0.3821** | | | | |
| DMS_RBD (original) | LucaVirus-Prot | | | | | | | | 0.9243 | 0.9470 |
| | LucaVirus-Mask | | | | | | | | 0.9277 | 0.9440 |
| | LucaVirus | | | | | | | | **0.9291** | **0.9477** |
| DMS_RBD (nucl) | LucaVirus-Nucl | | | | | | | | 0.8663 | 0.9193 |
| | LucaVirus-Mask | | | | | | | | 0.8660 | 0.9212 |
| | LucaVirus | | | | | | | | **0.8692** | **0.9247** |
| RdRP | LucaVirus-Prot | 0.9999 | 0.9981 | 0.9999 | 0.9999 | 0.9981 | | | | |
| | LucaVirus-Mask | 0.9997 | 0.9944 | 0.9998 | 0.9983 | 0.9943 | | | | |
| | LucaVirus | **1.0** | **1.0** | **1.0** | **1.0** | **1.0** | | | | |
| ViralCapsid | LucaVirus-Prot | **0.9988** | **0.9988** | **0.9997** | **0.9996** | **0.9976** | | | | |
| | LucaVirus-Mask | 0.9983 | 0.9983 | 0.9996 | 0.9992 | 0.9966 | | | | |
| | LucaVirus | 0.9985 | 0.9985 | 0.9996 | 0.9993 | 0.9971 | | | | |
| VirusEC4 | LucaVirus-Prot | | | | | | **0.9929** | **0.9965** | | |
| | LucaVirus-Mask | | | | | | 0.9822 | 0.9912 | | |
| | LucaVirus | | | | | | 0.9907 | 0.9957 | | |
| DMS_EVA | LucaVirus-Mask | | | | | | | | 0.7445 | 0.7820 |
| | LucaVirus-Prot | | | | | | | | 0.7562 | 0.7912 |
| | LucaVirus | | | | | | | | **0.7616** | **0.7984** |

Note: The antibody binding performance metrics in this table are based on all 98 antibodies, including those with weak binding affinity.

**Table S4. Performance comparison of LucaVirus and LucaOne on ten general tasks.**

| Task | Model | Acc | F1 | PR-AUC | Spearman |
|---|---|---|---|---|---|
| CentralDogma | LucaOne | **0.8453** | **0.7392** | **0.8617** | / |
| | LucaVirus | 0.7537 | 0.592758 | 0.680429 | / |
| SupKTax | LucaOne | 0.947 | 0.9467 | 0.9591 | / |
| | LucaVirus | **0.949** | **0.947768** | **0.959657** | / |
| GenusTax | LucaOne | 0.817 | 0.8069 | 0.8633 | / |
| | LucaVirus | **0.863** | **0.857885** | **0.901058** | / |
| SpeciesTax | LucaOne | 0.75 | 0.7345 | 0.7927 | / |
| | LucaVirus | **0.784** | **0.769504** | 0.782852 | / |
| ncRNAFam | LucaOne | **0.9864** | 0.9117 | **0.9622** | / |
| | LucaVirus | 0.982559 | 0.911888 | 0.952142 | / |
| ProtLoc | LucaOne | **0.9452** | **0.9378** | 0.9692 | / |
| | LucaVirus | 0.93634 | 0.93306 | **0.969878** | / |
| ProtStab | LucaOne | / | / | / | **0.7718** |
| | LucaVirus | / | / | / | 0.743177 |
| InfA | LucaOne | 1.0 | 1.0 | 1.0 | / |
| | LucaVirus | 1.0 | 1.0 | 1.0 | / |
| PPI | LucaOne | 0.9774 | 0.9799 | 0.9922 | / |
| | LucaVirus | **0.979394** | **0.981727** | **0.992441** | / |
| ncRPI | LucaOne | **0.9479** | **0.947** | 0.9781 | / |
| | LucaVirus | 0.947887 | 0.946801 | **0.978143** | / |

**Table S5. PERMANOVA analysis of embedding spaces for differentiating sequence features.** Statistical testing of model embeddings' capacity to separate sequences by three distinct biological features: amino acid identity, codon phase, and codon type.

| **Variable** | **R2** | **R2_Adj** | **P-value** | **Model_Name** |
|---|---|---|---|---|
| base | 0.20229846 | 0.20217878 | 0.001 | LucaOne |
| codon_phase | 0.04295975 | 0.04286403 | 0.001 | LucaOne |
| aminoacid | 0.10865857 | 0.10772163 | 0.001 | LucaOne |
| codon | 0.14390163 | 0.14115318 | 0.001 | LucaOne |
| Full_Model | 0.39108998 | 0.38833736 | 0.001 | LucaOne |
| base | 0.08336025 | 0.08322273 | 0.001 | LucaVirus |
| codon_phase | 0.02812825 | 0.02803104 | 0.001 | LucaVirus |
| aminoacid | 0.056764 | 0.05577251 | 0.001 | LucaVirus |
| codon | 0.07781359 | 0.07485297 | 0.001 | LucaVirus |
| Full_Model | 0.20462631 | 0.20103077 | 0.001 | LucaVirus |
| base | 0.11903172 | 0.11889955 | 0.001 | LucaVirus-Nucl |
| codon_phase | 0.03722265 | 0.03712636 | 0.001 | LucaVirus-Nucl |
| aminoacid | 0.07113074 | 0.07015435 | 0.001 | LucaVirus-Nucl |
| codon | 0.10059895 | 0.09771149 | 0.001 | LucaVirus-Nucl |
| Full_Model | 0.27487034 | 0.27159234 | 0.001 | LucaVirus-Nucl |
| base | 0.09953561 | 0.09940051 | 0.001 | LucaVirus-Mask |
| codon_phase | 0.02675398 | 0.02665664 | 0.001 | LucaVirus-Mask |
| aminoacid | 0.06487574 | 0.06389278 | 0.001 | LucaVirus-Mask |
| codon | 0.08852835 | 0.08560213 | 0.001 | LucaVirus-Mask |
| Full_Model | 0.22786513 | 0.22437464 | 0.001 | LucaVirus-Mask |
| base | 0.03261618 | 0.03247104 | 0.001 | EVO1-8K |
| codon_phase | 0.04218625 | 0.04209045 | 0.001 | EVO1-8K |
| aminoacid | 0.0560819 | 0.05508969 | 0.001 | EVO1-8K |
| codon | 0.07264765 | 0.06967044 | 0.001 | EVO1-8K |
| Full_Model | 0.16064847 | 0.15685412 | 0.001 | EVO1-8K |
| base | 0.03964316 | 0.03949908 | 0.001 | EVO2-7B |
| codon_phase | 0.03088893 | 0.030792 | 0.001 | EVO2-7B |
| aminoacid | 0.03546567 | 0.03445179 | 0.001 | EVO2-7B |
| codon | 0.05147731 | 0.04843214 | 0.001 | EVO2-7B |
| Full_Model | 0.13530725 | 0.13139835 | 0.001 | EVO2-7B |
| base | 0.00279747 | 0.00264786 | 0.001 | NT-V2-500M |
| codon_phase | 6.44E-05 | -3.56E-05 | 0.49 | NT-V2-500M |
| aminoacid | 0.00374634 | 0.00269913 | 0.001 | NT-V2-500M |
| codon | 0.01799032 | 0.01483764 | 0.001 | NT-V2-500M |
| Full_Model | 0.02111109 | 0.01668596 | 0.001 | NT-V2-500M |

**Table S6. Linear probing for genomic feature classification from model embeddings.** Reports classification performance (e.g., accuracy) of simple linear models trained to predict four distinct genomic features from frozen embeddings: CDS vs. non-CDS regions, amino acid identity, codon phase (reading frame), and codon type.

| | **Acc** | | | | | | **F1** | | | | | |
|---|---|---|---|---|---|---|---|---|---|---|---|---|
| **Model** | **CDS** | **base** | **phase** | **codon** | **aa** | **RBFS_phase** | **CDS** | **base** | **phase** | **codon** | **aa** | **RBFS_phase** |
| LucaVirus | 0.9380 | 1.0000 | 0.9163 | 0.8794 | 0.8642 | 0.7338 | 0.9380 | 1.0000 | 0.9163 | 0.8768 | 0.8616 | 0.7366 |
| LucaVirus-Nucl | 0.9330 | 1.0000 | 0.9113 | 0.8675 | 0.8437 | 0.7015 | 0.9330 | 1.0000 | 0.9113 | 0.8649 | 0.8409 | 0.6980 |
| LucaVirus-Mask | 0.9240 | 1.0000 | 0.9127 | 0.8811 | 0.8726 | 0.6915 | 0.9240 | 1.0000 | 0.9127 | 0.8782 | 0.8702 | 0.6919 |
| LucaOne | 0.9295 | 1.0000 | 0.9013 | 0.8637 | 0.8585 | 0.6642 | 0.9294 | 1.0000 | 0.9014 | 0.8596 | 0.8560 | 0.6642 |
| Evo-1-8k | 0.6285 | 0.3862 | 0.5580 | 0.0455 | 0.1020 | 0.4030 | 0.6284 | 0.3432 | 0.5394 | 0.0278 | 0.0784 | 0.4002 |
| Evo2-7B | 0.7425 | 0.9695 | 0.8033 | 0.1053 | 0.1863 | 0.6169 | 0.7402 | 0.9696 | 0.8041 | 0.0887 | 0.1509 | 0.6198 |
| NT-v2-500M | 0.7465 | 0.3098 | 0.3363 | 0.0384 | 0.0785 | 0.3234 | 0.7463 | 0.3081 | 0.3316 | 0.0322 | 0.0669 | 0.2891 |

**Table S7. Correlation between language model embeddings and genetic divergence of protein sequences.** Best metrics are colored red.

| | Metrics | Unseen Proteins | | | | RefSeq Proteins | | | |
|---|---|---|---|---|---|---|---|---|---|
| | | LucaVirus | LucaOne | ESM2 | ESMC | LucaVirus | LucaOne | ESM2 | ESMC |
| Pearson Correlation | Cosine distance vs. *p* distance | 0.71788789 | 0.67292459 | 0.46732339 | 0.46945701 | 0.73814079 | 0.86718098 | 0.75981986 | 0.77930478 |
| | *P* value | 0 | 0 | 0 | 0 | 0 | 0 | 0 | 0 |
| | Euclidean distance vs. *p* distance | 0.77199537 | 0.68346771 | 0.55707626 | 0.55494094 | 0.88842785 | 0.89616379 | 0.89246256 | 0.89877686 |
| | *P* value | 0 | 0 | 0 | 0 | 0 | 0 | 0 | 0 |
| | Cosine distance vs. phylogenetic distance* | 0.52350999 | 0.53489835 | 0.37349514 | 0.39681662 | 0.57705498 | 0.74245948 | 0.65126758 | 0.66326018 |
| | *P* value | 0 | 0 | 0 | 0 | 0 | 0 | 0 | 0 |
| | Euclidean distance vs. phylogenetic distance | 0.54526862 | 0.51891972 | 0.42307802 | 0.44204553 | 0.678809 | 0.71812782 | 0.71949099 | 0.72971626 |
| | *P* value | 0 | 0 | 0 | 0 | 0 | 0 | 0 | 0 |
| Spearman Correlation | Cosine distance vs. *p* distance | 0.69488777 | 0.69120009 | 0.53346517 | 0.52730391 | 0.71814951 | 0.86579372 | 0.86433049 | 0.86534489 |
| | *P* value | 0 | 0 | 0 | 0 | 0 | 0 | 0 | 0 |
| | Euclidean distance vs. *p* distance | 0.74883721 | 0.66831573 | 0.55415761 | 0.58649128 | 0.82507138 | 0.84648674 | 0.87746318 | 0.88903646 |
| | *P* value | 0 | 0 | 0 | 0 | 0 | 0 | 0 | 0 |
| | Cosine distance vs. phylogenetic distance | 0.52966593 | 0.56079703 | 0.41000241 | 0.41266814 | 0.70513822 | 0.849783 | 0.82559709 | 0.82786262 |
| | *P* value | 0 | 0 | 0 | 0 | 0 | 0 | 0 | 0 |
| | Euclidean distance vs. phylogenetic distance | 0.57467934 | 0.5362321 | 0.42339089 | 0.45910491 | 0.80493673 | 0.8389194 | 0.8387283 | 0.85415401 |
| | *P* value | 0 | 0 | 0 | 0 | 0 | 0 | 0 | 0 |

*Phylogenetic distances were computed from a maximum likelihood (ML) phylogenetic tree and represent the estimated number of substitutions per site between pairs of homologous sequences.

**Table S8. Correlation between language model embeddings and genetic divergence of nucleotide sequences.** Best metrics are colored red.

| | Metrics | Unseen Genes (nucleotide) | | RefSeq Genes (nucleotide) | |
|---|---|---|---|---|---|
| | | **LucaVirus** | **LucaOne** | **LucaVirus** | **LucaOne** |
| Pearson Correlation | Cosine distance vs. *p* distance | 0.76357767 | 0.64819183 | 0.90289818 | 0.77512818 |
| | *P* value | 0 | 0 | 0 | 0 |
| | Euclidean distance vs. *p* distance | 0.82627862 | 0.71496931 | 0.93553203 | 0.90442508 |
| | *P* value | 0 | 0 | 0 | 0 |
| | Cosine distance vs. phylogenetic distance* | 0.53775195 | 0.48494253 | 0.67405342 | 0.56478566 |
| | *P* value | 0 | 0 | 0 | 0 |
| | Euclidean distance vs. phylogenetic distance | 0.56996224 | 0.51259255 | 0.67548841 | 0.63976091 |
| | *P* value | 0 | 0 | 0 | 0 |
| Spearman Correlation | Cosine distance vs. *p* distance | 0.72093615 | 0.6149491 | 0.84638142 | 0.81946972 |
| | *P* value | 0 | 0 | 0 | 0 |
| | Euclidean distance vs. *p* distance | 0.78376811 | 0.64234757 | 0.89161081 | 0.83500575 |
| | *P* value | 0 | 0 | 0 | 0 |
| | Cosine distance vs. phylogenetic distance | 0.55254023 | 0.48869635 | 0.8005068 | 0.76488604 |
| | *P* value | 0 | 0 | 0 | 0 |
| | Euclidean distance vs. phylogenetic distance | 0.61729172 | 0.51256412 | 0.8488809 | 0.78273279 |
| | *P* value | 0 | 0 | 0 | 0 |

*Phylogenetic distances were computed from a maximum likelihood (ML) phylogenetic tree and represent the estimated number of substitutions per site between pairs of homologous sequences.

**Table S9. Performance comparison of different LLMs on classification tasks.**

| Tasks | Method | Acc | F1 | AUC | PR-AUC | MCC |
|---|---|---|---|---|---|---|
| DeepAbBindv2_genome | ESM2+DNABert2 | 0.733333 | 0.529412 | 0.665801 | 0.580468 | 0.371831 |
| | ESM2+NT | 0.716667 | 0.548673 | 0.672499 | **0.608799** | 0.348529 |
| | LucaOne | 0.733333 | 0.555556 | **0.702023** | 0.606699 | 0.37943 |
| | LucaVirus | **0.738889** | **0.576577** | 0.691361 | 0.552956 | **0.397208** |
| DeepAbBindv2_nucl | ESM2+DNABert2 | 0.677778 | 0.462963 | 0.658488 | 0.509923 | 0.245408 |
| | ESM2+NT | 0.672222 | 0.504202 | 0.656985 | 0.53887 | 0.260546 |
| | LucaOne | 0.694444 | 0.495413 | 0.642701 | 0.492226 | 0.287761 |
| | LucaVirus | **0.733333** | **0.538462** | **0.695052** | **0.60243** | **0.37409** |
| DeepAbBindv2_original | ESMC | 0.688889 | 0.517241 | 0.663546 | 0.601243 | 0.290843 |
| | ESM2 | 0.7 | 0.517857 | 0.683024 | 0.560254 | 0.307427 |
| | LucaOne | 0.705556 | 0.522523 | 0.687534 | 0.550662 | 0.318408 |
| | LucaVirus | **0.727778** | **0.581197** | **0.712411** | **0.610665** | **0.382117** |
| Phage_PVP | ESMC | 0.989348 | 0.953226 | 0.995852 | 0.963023 | 0.978936 |
| | ESM2 | **0.991209** | **0.958569** | 0.995687 | **0.965605** | **0.982621** |
| | LucaOne | 0.987238 | 0.945649 | **0.996022** | 0.953041 | 0.974751 |
| | LucaVirus | 0.987176 | 0.946065 | 0.994515 | 0.947646 | 0.974588 |
| RdRP | ESMC | 0.999865 | 0.99723 | 0.999999 | 0.99998 | 0.997165 |
| | ESM2 | 0.999865 | 0.99723 | 0.999999 | 0.999976 | 0.997165 |
| | LucaOne | 0.99991 | 0.998145 | 0.999997 | 0.999884 | 0.998101 |
| | LucaVirus | **1.0** | **1.0** | **1.0** | **1.0** | **1.0** |
| ViralCapsid | ESMC | 0.997739 | 0.99774 | 0.999488 | 0.998993 | 0.995479 |
| | ESM2 | 0.998308 | 0.998308 | 0.999639 | 0.999299 | 0.996617 |
| | LucaOne | 0.998293 | 0.998293 | 0.999646 | 0.999341 | 0.996586 |
| | LucaVirus | **0.998554** | **0.998554** | **0.999656** | **0.999371** | **0.997109** |

**Table S10. Performance comparison of different LLMs on multi-label classification task.**

| Tasks | Method | Jaccard | fmax |
|---|---|---|---|
| VirusEC4 | ESMC | 0.990217 | 0.995257 |
| | ESM2 | **0.993335** | **0.996788** |
| | LucaOne | 0.991878 | 0.99632 |
| | LucaVirus | 0.990781 | 0.995724 |

**Table S11. Performance comparison of different LLMs on regression tasks.**

| Tasks | Method | SRCC (Spearman's correlation coefficient) | PRCC (Pearson correlation coefficient) |
|---|---|---|---|
| DMS_EVA | ESMC-600M | 0.683563 | 0.720946 |
| | ESM2-3B | 0.758408 | 0.791698 |
| | LucaOne | 0.753631 | 0.785300 |
| | LucaVirus | **0.761608** | **0.798425** |
| DMS_Bind_Reps_Strain | ESMC-600M | 0.706095 | 0.769857 |
| | ESM2-3B | 0.837381 | 0.887300 |
| | LucaOne | 0.901307 | 0.943520 |
| | LucaVirus | **0.929127** | **0.947747** |
| DMS_Bind_Reps_Strain_Nucl | DNABert2 | 0.794032 | 0.870466 |
| | NT-2.5B-MultiSpecies | 0.846765 | 0.909852 |
| | LucaOne | 0.856909 | 0.924048 |
| | LucaVirus | **0.869273** | **0.924707** |
| HK19_immune_escape_per_serum | ESMC-600M | 0.239878 | 0.231521 |
| | ESM2-3B | 0.512224 | 0.612066 |
| | LucaOne | 0.502234 | 0.642996 |
| | LucaVirus | **0.537990** | **0.723446** |

**Table S12. Confusion matrix of the wet-lab validated antibodies prediction by LucaVirus model (weak binding (+/++) excluded).**

| **Variant: Wuhan-Hu-1** | | Prediction | |
|---|---|---|---|
| | | Neg. | Pos. |
| Ground Truth | Neg. | 53 | 6 |
| | Pos. | 6 | 13 |
| **Variant: Omicron** |  | Prediction |  |
| | | Neg. | Pos. |
| Ground Truth | Neg. | 71 | 4 |
| | Pos. | 3 | 7 |

**Table S13. Wet-lab validated antibodies information.**

| Name | Published Name | cell_id | WT Spike binding | Omicron Spike binding |
|---|---|---|---|---|
| I12 | CAV-C12 | I3_GTAGTCAAGGTGCAAC_1 | + | + |
| I14-2 | CAV-C14 | I4_CGTCAGGAGTGACTCT_1 | + | - |
| I17 | CAV-C17 | I1_CGGAGTCAGAAGGGTA_1 | + | - |
| I23 | CAV-C23 | I5_TTTCCTCCATAGACTC_1 | - | - |
| I28 | CAV-C28 | I5_GGATTACTCCTAGGGC_1 | - | - |
| I31 | CAV-C31 | I5_TCAGCTCTCGCTTAGA_1 | - | - |
| I35 | CAV-C35 | I1_CGGACTGGTATTAGCC_1 | - | - |
| I39 | CAV-C39 | SCoV1-M0_AGCTTGAAGGAGTTTA_1 | +++ | - |
| I40 | CAV-C40 | SCoV13-M0_AAACGGGTCTGCGGCA_1 | + | + |
| I44 | CAV-C44 | K1_CCACTACGTCTCTTTA_1 | +++ | +++ |
| I47 | CAV-C47 | K1_CTTAGGAGTTTAGCTG_1 | +++ | ++ |
| I59 | CAV-C59 | I7_GAACATCCAAGCGATG_1 | + | - |
| I60 | CAV-C60 | I8_ACGCCAGTCCCTAATT_1 | - | - |
| I61 | CAV-C61 | I8_TCAATCTGTGTGGCTC_1 | +++ | +++ |
| I65 | CAV-C65 | I7_CGCTGGAGTCGAAAGC_1 | +++ | + |
| I68 | CAV-C68 | I9_AACCGCGTCTGCAGTA_1 | +++ | +++ |
| I74 | CAV-C74 | I10_TGCTGCTGTATATCCG_1 | +++ | +++ |
| I75 | CAV-C75 | I10_GCGCGATAGCGAAGGG_1 | +++ | +++ |
| I76 | CAV-C76 | I9_CTACACCTCGAACGGA_1 | +++ | +++ |
| I77 | CAV-C77 | I9_GTCGTAAAGACCTAGG_1 | +++ | ++ |
| I79 | CAV-C79 | I10_CACACTCGTGAGGGAG_1 | +++ | +++ |
| I90 | CAV-C90 | SCoV11-M0_GACACGCTCCTAGAAC_1 | + | - |
| I91 | CAV-C91 | SCoV1-M0_TGATTTCTCATCGCTC_1 | - | - |
| I95 | CAV-C95 | I1_GGATTACCAAGCTGGA_1 | ++ | + |
| I96 | CAV-C96 | I2_GTACTCCCAGCCTTGG_1 | ++ | - |
| I97 | CAV-C97 | I3_TGCGGGTGTGTAAGTA_1 | ++ | + |
| I98 | CAV-C98 | I7_CACAGGCTCTTAGAGC_1 | + | - |
| CD26 | CAV-C26 | DG15-P1-B2 | + | - |
| CD155 | CAV-C155 | SG04-P1-H9 | + | - |
| CD217 | CAV-C217 | DG18-P2-D7 | +++ | - |
| CD218 | CAV-C218 | DG18-P2-E4 | +++ | - |
| CD219 | CAV-C219 | SG04-P2-A11 | + | + |
| CD236 | CAV-C236 | SC50-P1-A5 | +++ | +++ |
| CD241 | CAV-C241 | WH46-C1 | + | + |

| | | | | |
|---|---|---|---|---|
| CD286 | CAV-C286 | WH19-P1-C4 | +++ | - |
| CD311 | CAV-C311 | DG29-P2-D6 | +++ | ++ |
| CD327 | CAV-C327 | SC2-074-P3-H3 | +++ | +++ |
| CD329 | CAV-C329 | SC2-074-P3-C4 | + | - |
| CD339 | CAV-C339 | WH28-P1-F5 | +++ | +++ |
| CD354 | CAV-C354 | WH2-1B-P3P5-C2 | ++ | - |
| CD355 | CAV-C355 | WH2-1B-P3P5-E11 | + | + |
| CD358 | CAV-C358 | WH2-1B-P3P5-D7 | - | - |
| CD362 | CAV-C362 | WH2-14-P2-A6 | + | - |
| CD368 | CAV-C368 | WH2-14-P2-E3 | + | - |
| CD380 | CAV-C380 | WH2-23-PC-A4 | ++ | + |
| CD385 | CAV-C385 | WH35-P2-H3 | +++ | ++ |
| I14-1 | | I4_CGTCAGGAGTGACTCT_1 | - | - |
| I20 | | K2_ACCGTAACAACGCACC_1 | - | - |
| I21 | | K1_TAGGCATCATGGGACA_1 | - | - |
| I22 | | K2_AGCGGTCAGTAACCCT_1 | - | - |
| I26 | | I2_CAGATCAAGGGCATGT_1 | - | - |
| I27 | | I5_GTCATTTTCATGCAAC_1 | - | - |
| I29 | | K1_CACAGGCAGTCCCACG_1 | - | - |
| I30 | | I4_CAGCCGAGTGTGAAAT_1 | - | - |
| I32 | | K1_CTAACTTCACCTGGTG_1 | - | - |
| I33 | | I6_AACGTTGTCCGCAAGC_1 | - | - |
| I34 | | I3_CAAGGCCTCGTCTGCT_1 | - | - |
| I37 | | I4_GCAGTTATCCATGCTC_1 | - | - |
| I38 | | SCoV1-M0_TCTATTGTCAGTACGT_1 | - | - |
| I41 | | I3_GATCGATAGGAGTACC_1 | - | - |
| I42 | | I3_TCAGGATCAGTCAGCC_1 | - | - |
| I43 | | I3_CATGACATCTCCGGTT_1 | - | - |
| I45 | | K2_GGTGAAGGTGTATGGG_1 | - | - |
| I46 | | K2_GTGTGCGCAGTGGGAT_1 | - | - |
| I48 | | I4_CGAGAAGCACGGCGTT_1 | - | - |
| I49 | | I5_CCTAGCTCAAGGCTCC_1 | - | - |
| I50 | | I6_AACCATGCACACCGCA_1 | - | - |
| I51 | | I3_GTTTCTATCCAAATGC_1 | - | - |
| I52 | | I5_ACACCCTTCCACTCCA_1 | - | - |
| I53 | | K2_ATTACTCAGATGTCGG_1 | - | - |
| I55 | | K1_GATGAAAGTGATAAAC_1 | - | - |
| I56 | | I7_GCAGCCACATTCCTGC_1 | - | - |
| I57 | | K4_TACCTTAGTTCACCTC_1 | - | - |

| | | | | |
|---|---|---|---|---|
| I58 | | I7_AAACGGGGTACCGAGA_1 | - | - |
| I63 | | K4_TGGCCAGCAAGGACAC_1 | - | - |
| I64 | | K4_AGCATACCATGACGGA_1 | - | - |
| I66 | | I7_TTGGCAAAGTTAGCGG_1 | - | - |
| I67 | | I8_CCTATTACAATCTGCA_1 | - | - |
| I69 | | I9_AACCGCGCATCGATTG_1 | - | - |
| I70 | | I9_CTACCCAAGCCTTGAT_1 | - | - |
| I71 | | I9_CTGATAGGTTAGTGGG_1 | - | - |
| I72 | | I9_CAGCTAAGTAACGCGA_1 | - | - |
| I73 | | I10_CTGCTGTTCCGAAGAG_1 | - | - |
| I78 | | I9_AGAGCTTTCACTGGGC_1 | - | - |
| I80 | | I9_CTACCCAGTACCGCTG_1 | - | - |
| I81 | | I9_TCAGCAATCATTCACT_1 | - | - |
| I82 | | I9_CCTACCAGTCTACCTC_1 | - | - |
| I83 | | I9_AGTGGGAGTGCCTTGG_1 | - | - |
| I84 | | I9_AGTAGTCTCGTACCGG_1 | - | - |
| I85 | | I10_CATCGGGTCCTAGTGA_1 | - | - |
| I86 | | I10_TCCCGATTCAGAGCTT_1 | - | - |
| I87 | | I10_ATCCACCCACGTCTCT_1 | - | - |
| I88 | | I10_TAAGCGTCAGGTGGAT_1 | - | - |
| I89 | | SCoV11-M0_CATATTCGTAGGAGTC_1 | - | - |
| I92 | | I6_CGCTTCAGTGTGGCTC_1 | - | - |
| I94 | | SCoV1-M0_TCCACACAGGCATGTG_1 | - | - |
| I99 | | I7_ATGCGATGTCTTTCAT_1 | - | - |
| I100 | | I10_GGTGAAGCACTGTGTA_1 | - | - |
